# Gene expression evolution is predicted by stronger indirect selection at more pleiotropic genes

**DOI:** 10.1101/2024.07.22.604294

**Authors:** Eva L. Koch, Charles Rocabert, Champak Beeravolu Reddy, Frédéric Guillaume

## Abstract

Changes in gene expression play a fundamental role in the process of adaptation and can provide insight into the genetic basis of adaptation. We utilized transcriptome-wide variation in gene expression as a means to uncover genes under selection for expression changes during adaptation to heat and drought stress, and to understand the nature of selection on gene expression traits of the red flour beetle *Tribolium castaneum*. We showed that estimates of genetic selection on transcript abundance were predictive of evolutionary changes in gene expression after 20 generations of adaptation in seven independent experimental lines. Having measured the genetic covariance between gene expression and relative fitness and among expression traits, we showed that evolutionary changes were driven more by indirect selection acting on genetically correlated partners rather than by direct selection acting on isolated genes. Consequently, genes with central positions in gene co-expression networks experienced stronger selection and exhibited larger evolutionary changes in expression. Our genomic analysis revealed that selection on expression levels is associated with parallel allele frequency changes (AFCs) in the respective genes. More pleiotropic genes and those carrying expression quantitative trait loci (eQTLs) showed a higher degree of parallel evolution. More generally, the stronger the parallelism of AFCs in a gene, the stronger its genetic selection. Contrary to previous evidence of constrained evolution at more connected genes, adaptation was driven by selection acting disproportionately on genes central to co-expression gene networks. We demonstrated that measures of selection at the transcriptome level can provide accurate evolutionary predictions and critical information on the molecular basis of rapid adaptation.

## Introduction

Understanding the genetic basis of adaptation is a central goal in evolutionary biology. Molecular studies of trait variation have highlighted the key role of gene expression in phenotypic variation. Although the importance of gene expression in adaptation is largely accepted (Fay and Wittkopp, 2008) and documented (*e.g*., Whitehead and Crawford, 2006; Fraser et al., 2010; Pritchard et al., 2017; Mack et al., 2018; Kelly et al., 2023), it remains difficult to understand how variation in thousands of expression levels translates into fitness variation and evolves in a coordinated manner. Many genes do not act in isolation, but are integrated into pathways, jointly contribute to the same functional processes, and are connected by various interactions. Thus, understanding how variation in the expression of individual genes is linked to fitness and how they may respond to selection remains challenging. Only a few recent studies have successfully connected gene expression variation with fitness variation in insects (McGuigan et al., 2014; Koch and Guillaume, 2020a), plants (Groen et al., 2020; Henry and Stinchcombe, 2025), or vertebrates (Ahmad et al., 2021). This approach has been used to identify genes whose expression variation is most important for fitness under specific stresses (*e.g*., drought in rice; Groen et al., 2020). However, we still have limited insight into the mechanisms underlying expression evolution and the extent to which it is predictable.

Evolutionary changes can only occur if at least some of the observed variation is caused by underlying genetic variation and exhibits genetic covariation with fitness (known as the Robertson-Price identity; Robertson, 1966; Price, 1970). It has been shown that gene expression levels can be considered quantitative traits, for which genetic variance and their genetic covariance with fitness can be estimated (Cheung and Spielman, 2002). While estimating this covariance can predict future changes, the underlying mechanisms remain unclear. Since expression levels are expected to be highly interconnected (Ayroles et al., 2009; McGraw et al., 2011; Blows et al., 2015), direct selection acting on one expression level leads to indirect selection on its correlated partners. Evolutionary changes might therefore occur because an expression level is directly targeted by selection, or because it is genetically correlated with other expression levels under selection, or both. Disentangling the effects of direct and indirect selection, though challenging, would provide detailed insights into the mechanisms by which certain expression levels change, their role in adaptation, and potential constraints. Key to this is the estimation of genetic covariances between individual expression levels.

The presence of genetic covariance among expression levels (Ayroles et al., 2009; McGraw et al., 2011; Blows et al., 2015) also implies that they cannot evolve independently, as selection acting on one of them would indirectly affect all correlated expression levels, potentially limiting responses to certain combinations of selective pressures. Genetic covariance is therefore often discussed in the context of evolutionary constraints and evolvability (Blows and Hoffmann, 2005; Walsh and Blows, 2009; Hansen, 2003; Hansen and Pelabon, 2021). In this context, pleiotropy—the property of genes affecting multiple traits—plays a key role by generating genetic and phenotypic covariance (Lande, 1980; Walsh and Blows, 2009). Pleiotropic genes are often considered to be under the strongest constraints, as any change is less likely to benefit all the traits they influence (Fisher, 1930; Orr, 2000). In gene expression, regulatory genes at central positions within regulatory networks controlling other genes’ expression are expected to be the most pleiotropic and, as a result, the most evolutionarily constrained (Stern and Orgogozo, 2008; Josephs et al., 2017).

On the other hand, genetic covariance can enhance selection responses and accelerate evolution when selection favours correlated responses. In this case, a single expression level experiences not only the direct effect of selection but also the combined effects of selection acting on all its genetically correlated partners, enhancing its response in the direction of selection (Lande, 1979; Agrawal and Stinchcombe, 2009). The evolutionary importance and potential constraints associated with covariances between expression levels remain largely unknown. While some studies found evidence for constraints (Fraser et al., 2002; Innocenti and Chenoweth, 2013; Josephs et al., 2015; Kita et al., 2017), other studies suggest that these effects might be weak (Steige et al., 2017; Glassberg et al., 2019) and rapid evolution of gene expression during adaptation to new environments has been demonstrated in several species (Yampolsky et al., 2012; Ghalambor et al., 2015; Huang and Agrawal, 2016; Koch and Guillaume, 2020b). Emerging evidence from experimental evolution experiments and studies of the genetic basis of local adaptation to environmental variation suggests that pleiotropy (or gene connectivity in gene networks; He and Zhang, 2006) plays a key positive rather than constraining role in adaptation (Rennison and Peichel, 2022; Ruelens et al., 2023; Thorhölludottir et al., 2023; Whiting et al., 2024). All in all, the potential for the evolution of gene expression is likely large but may strongly depends on the regulatory and functional network context of expression variation (Jordan et al., 2004; Innocenti and Chenoweth, 2013; Collet et al., 2018).

A powerful framework for analysing expression and its underlying network lies in identifying coexpression modules, groups of genes whose expression levels covary across in-dividuals (Langfelder and Horvath, 2008). Although based on phenotypic correlations rather than causal relationships, modules can help reveal functionally related gene sets and highlight the most connected genes within networks that may act as key regulators (Jeong et al., 2001; Liseron-Monfils and Ware, 2015; Vengatharajuloo et al., 2023). Moreover, examining whether modules are preserved or reorganised under environmental stress can identify the processes most affected and may help uncover mechanisms underlying evolutionary adaptation (*e.g*., des Marais et al., 2017; Helsen et al., 2019).

Evolutionary changes in gene expression must be accompanied by corresponding alterations at the genomic level. Studying allele frequency changes during adaptation and their parallelism across replicated populations or experimental lines can help us gain a more comprehensive understanding of the genetic basis of adaption and of gene expression evolution (Barghi et al., 2020). Studying parallel evolution across replicates is commonly used to address the challenge of distinguishing between changes caused by genetic drift and those driven by selection, and conclude for a predominant role of selection (Yeaman et al., 2018; Barghi et al., 2019; Whiting et al., 2024). Furthermore, identifying causal genomic variation underlying expression changes with expression QTL (eQTL) mapping can deepen our understanding of the genetic architecture of gene expression and how selection on expression levels leads to evolutionary changes (Fraser et al., 2010; des Marais et al., 2013; Josephs et al., 2015; Kelly et al., 2023).

Here, we took advantage of individual transcriptome (RNA-seq) and fitness data from two time points measured in 486 females within replicated experimental evolution lines of the model organism *Tribolium castaneum* (red flour beetle) adapting to a hot-dry environment. Previous results from this experiment demonstrated that this stressful environment strongly reduces average fitness, as measured by the number of surviving offspring per female (Koch and Guillaume, 2020a), and that adaptation over 20 generations led to significant evolutionary changes in gene expression (Koch and Guillaume, 2020b). Constructing gene co-expression networks allowed us to explore how network structure changes under stress. Combining this with selection estimates provided insights into how selection acted on different network components and how this related to improving fitness under stressful conditions. Using experimental evolution, we could then investigate how expression changed during adaptation, contrast these outcomes with our predictions made in the first generation and explore associated changes at the genomic level.

Our goals were: 1) to evaluate the predictability of gene expression evolution by assessing whether evolutionary changes were consistent with predictions made in the first generation; 2) and to gain a detailed understanding of the causes of evolutionary changes in gene expression and their underlying mechanisms. To do so, we quantified and disentangled the contributions of direct and indirect selection to the predicted evolutionary change (Stinchcombe et al., 2014). Using this approach enabled us to address crucial questions regarding the genetic architecture of gene expression adaptation: Are there a few genes that are direct targets of selection, with the majority of genes showing a correlated response? Is selection acting on many gene expression levels simultaneously, and can position in gene expression networks predict their response to selection? 3) Finally, we aimed to link evolutionary changes in expression levels with genetic changes. Having seven replicate selection lines, we tested for parallel allele frequency changes (AFCs) and whether these changes were associated with genes under selection for expression changes. Furthermore, we tested whether SNPs associated with variation in expression patterns (eQTLs) showed parallel AFCs in the different selection lines.

## Material and Methods

Full details of animal rearing, setting up the paternal half-sib/full-sib crosses, fitness assays, experimental evolution and testing for adaptation can be found in Koch and Guillaume (2020a,b). Underlying data and result files (referenced as Data S1-S10) are available at Zenodo DOI: 10.5281/zenodo.15298510.

### Crossing for estimating genetic variances

We estimated heritable genetic variation (*V*_*A*_) of transcriptome-wide gene expression levels and individual fitness (number of surviving offspring per individual), and their genetic covariance in the first generation of our experiment from combined measurements of gene expression (RNA-seq) and fitness for 486 females obtained from a paternal half-sib/full-sib breeding design, which provides an unbiased estimate of *V*_*A*_ (Lynch and Walsh, 1998) (Figure 1). Our experimental lines were derived from the *Tribolium castaneum* Cro1 strain (Milutinović et al., 2013), which had been collected from a wild population in 2010 and adapted to lab standard conditions (33°C, 70% relative humidity—or r.H.) for more than 20 generations. To estimate genetic variances in Control and Hot-Dry (37°C, 30% r.H.), hereafter referred to as CT and HD, respectively, we performed a paternal half-sib breeding design with 147 half-sib families (see Koch and Guillaume, 2020a; Koch et al., 2020). Half-sib families were created by mating a virgin male to three virgin females. Half- and full-sib families were then split across conditions at the egg stage.

**Figure 1.**
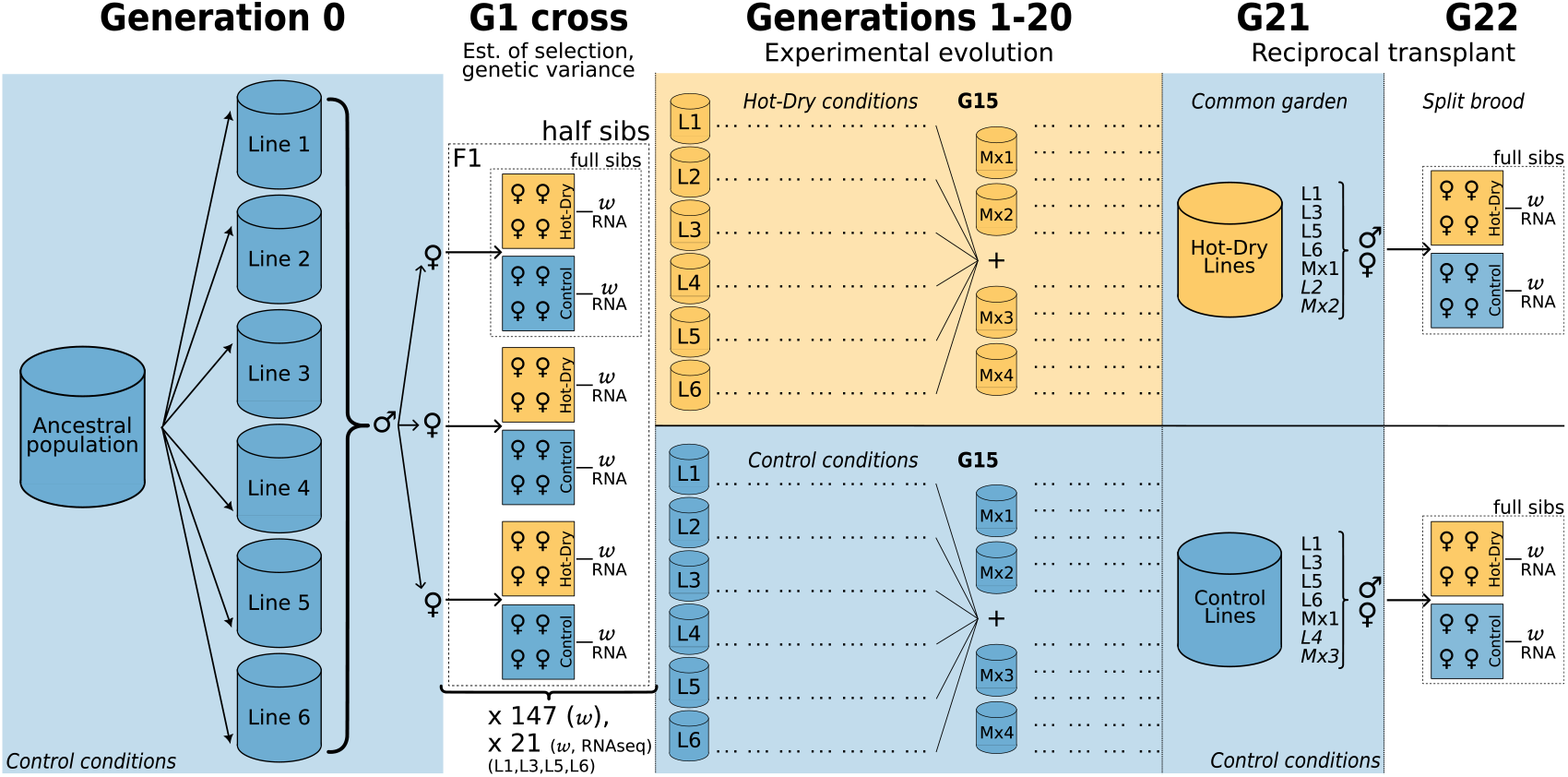
Experimental design. Six experimental lines each containing 120 adults beetles with equal sex ratio were derived from a large ancestral population adapted to Control conditions (30°C, 70% relative humidity). A half-sib/full-sib cross with 147 half-sib families was performed from the first brood of Generation 0. The F1 of the cross was raised in Control and Hot-Dry (37°C, 30% r.H.) conditions from egg to adult. The F1’s fitness (*w*) and RNA were measured (see main text). The experimental lines 1-6 were replicated in Control and Hot-Dry conditions and propagated for 20 generations with a constant adult size of 120 (breeders). At generation 15, four Mixed lines were added to the design. They were created by mixing 20 adults from each line. At G20, the Hot-Dry and Control lines were brought to a common garden in Control. The brood of G20 was raised for one full generation (egg to adults) in Control. The G21 adults were then used to perform a reciprocal transplant (G22) by splitting their brood in Control and Hot-Dry conditions. The G22’s fitness and RNA were measured (see main text).

We estimated heritable genetic variance for fitness (*V*_*A,w*_) from the complete set of 147 half-sib families created during the crossing experiment (*ca*. 3000 individuals). *V*_*A*_ for gene expression was estimated from a subset of female individuals from which we measured both fitness and gene expression levels in 21 half-sib families (486 females). Fitness was measured as the reproductive output per female of 2,937 virgin females (1,518 in Control and 1,419 in Hot-Dry) that had been mated with an unrelated male from the same condition.

We previously showed that average fitness decreased from 49.1 in CT to 25.7 in HD. Yet, the heritable genetic variance of relative fitness 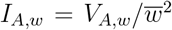 was higher in HD than CT (percent increase of mean fitness in HD = 3.2%, and in CT = 1.6%, see Koch et al., 2020), pointing at a higher adaptive potential in the stressful environment than in the control (Houle, 1992; Lynch and Walsh, 1998).

### Experimental evolution and reciprocal transplant

We used six replicate lines per condition originating from the Cro1 ancestral population, which was used for the fitness assay in G1 (see above). We let each line adapt for 20 non-overlapping generations in CT and HD. Each line consisted of 120 breeding individuals. Additional “mixed” lines were created at generation 15 by taking 20 individuals of each selection line and placing them into a new vial. In total we had ten selection lines per condition: six original lines and four mixed lines.

We assessed adaptation by testing for significant differences in mean fitness in HD conditions between females originating from CT and HD lines. After one full generation in CT conditions (generation 21; Figure 1, see also Koch and Guillaume, 2020b), full-sib families were created within each selection line and then split across CT and HD conditions in the egg stage (generation 22; Figure 1). With this design, we could assess fitness differences of full-sib families in both conditions from lines originating from either CT or HD. Fitness of G22 females was assessed by mating a virgin male and a virgin female of the same selection line, but from different families and counting the number of adult offspring. After four days of mating and egg laying, males and females were removed from the medium and flash-frozen in liquid nitrogen to be stored at −80°C until RNA extraction and sequencing. Samples were always frozen at the same time of the day (morning) to minimize circadian rhythm effects.

The transplant revealed that HD lines had higher average fitness (30.3) than CT lines in HD (19.5) and had thus adapted to HD conditions after 20 generations (Koch and Guillaume, 2020b).

### Sampling, RNA extraction, library preparation, sequencing and data processing

We chose whole-body RNA sequencing to gain a comprehensive systemic view of gene expression changes in response to environmental conditions. While shifts in body composition (*e.g*., organ size changes) may contribute to differential expression patterns, these are integral aspects of organismal adaptation and provide valuable biological insights. Our approach ensures that we capture all relevant transcriptional changes, allowing for a systemic understanding of the evolutionary responses to selection without making any preselection regarding the organ or tissue most relevant for adaptation. RNA-seq (whole transcript sequencing) was performed on a total of 644 samples divided as 486 female beetles with known fitness at generation 1 (244 from CT and 242 from HD) and 158 females with known fitness at generation 22 (90 from CT and 68 from HD). Generation 1 samples originated from a half-sib/full-sib cross (see above) with approximately 21 sires, 3 dams/sires and 8 full-sib females/dam, which were separately grown in CT and HD in batch of 4 females per full-sib family. In generation 22, we sampled four females per family per environment from seven full-sib families and lines (details in *Allelic frequency changes* subsection). Since the original experiment consisted in two additional treatments (Dry and Hot, see Koch and Guillaume, 2020a, Koch and Guillaume, 2020b), the total number of RNAseq samples amounts to 822 individuals. The extra 178 transcriptomes originate from Hot and Dry (93 in G1, 85 in G22) and did not enter our quantitative genetics analyses but partic-ipated in the overall detection of genetic variants from nucleotides polymorphisms (see next section). Of the 486 G1 females, 90 were analysed in Koch and Guillaume (2020a). The G22 samples were analysed in Koch and Guillaume (2020b) to test for evolutionary changes in gene expression. This analysis yielded 55 genes with a significant evolutionary response of their transcript abundance in HD at generation 22 after correction for multiple testing (Koch and Guillaume, 2020b). From the same study, we retrieved the set of up- and down-regulated genes in HD lines compared to their counterparts in CT lines in HD.

The same extraction and library preparation protocols were followed for each set of individuals sequenced separately at generation 1 and 22 in all treatments. For full details see Koch and Guillaume, 2020a,b. The treatments were randomized during RNA-extraction, library preparation (selecting mRNA by poly(A) selection), and sequencing. Single-end sequencing was performed in five runs (36 libraries pooled for each run) on the Illumina NextSeq 500 (Illumina, Inc, California, USA) using the 75 cycles (= read length 75bp) High Output Kit. Each run resulted in 550-600 million reads that passed the internal sequencer filter. After quality control using FastQC (www.bioinformatics.bbsrc.ac.uk/projects/fastqc), reads were mapped against the reference genome (ftp://ftp.ensemblgenomes.org/pub/release30/metazoa/gtf/tribolium_castaneum/Tribolium_castaneum.Tcas3.30.gtf.gz) with STAR v.2.5 (Dobin et al. 2013; adapters were trimmed and the first 10 bases were hard trimmed, minimum average quality Q10, minimum tail quality 10, minimum read length 20). We then used FeatureCounts (Liao et al., 2014) to count the number of reads that mapped to each gene in the reference genome. Mapping as well as read counting was performed within the data analysis framework SUSHI (Hatakeyama et al., 2016).

### Quantitative Genetics analyses of heritable variation and selection on gene expression

We estimated genetic selection acting on gene expression from the genetic covariance of the trait with relative fitness *s*_*g*_ = *cov*_*A*_(*z, w*) (Robertson, 1966; Price, 1970) in 486 individuals sampled from the cross performed at the first generation of the experiment, with 244 from CT and 242 from HD. We implemented a Bayesian sparse factor analysis of genetic covariance matrices (BSFG) method as described in Runcie and Mukherjee (2013). BSFG uses a linear mixed model and sparse factorization to estimate the additive genetic covariance matrix **G** and the residual covariance matrix **R** among traits. We utilized BSFG to estimate the genetic covariance between the expression level of each gene and relative fitness and between pairs of genes. To run BSFG, we used the MATLAB (MATLAB version 2014b) code provided at www2.stat.duke.edu/∼sayan/bfgr/Instructions/soft.html. We included a batch effect (sequencing runs) as fixed factor and mother identity as random factor to account for potential maternal and common environment effects. Since it was not possible to analyze the whole data in one single analysis due to computational limitations, we divided the whole data set randomly into ten smaller subsets (each consisting of 1181 genes in CT, 1189 genes in HD). We repeated this subdivision five times, resulting in five subdivisions each consisting of ten different subsets. Relative fitness was included as a trait in all subsets. All of these subsets were analyzed three times (three replicates) with 50,000 iterations of BSFG each. The first 20,000 iterations were discarded as burn-in and the remaining chain thinned for every 30th sample resulting in 1000 posterior samples for each replicate. Convergence was assessed by using Gelman-Rubinstein diagnostics 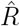 (Gelman and Rubin, 2007; Brooks and Gelman, 1998) for the three replicates. MATLAB output was further analyzed in R (version 3.4.2, R Core Team, 2017). The posterior distribution of **G** was calculated from the output of BSFG as described in Runcie and Mukherjee (2013). The genetic correlation of transcript expression with fitness (*r*_*g*_) was computed as 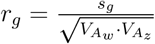, where 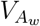 and 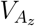 are the additive genetic variances of relative fitness and of the transcript expression level, respectively. Additive genetic variances correspond to the diagonal elements of **G**. The off-diagonal elements hold the pairwise genetic covariance between gene expression levels. We further decomposed genetic selection *s*_*g*_ into its direct and indirect selection components following Stinchcombe et al. (2014). Accordingly, we computed the gradient of direct genetic selection on gene expression levels as 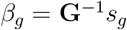 from the posterior distributions of **G** and *s*_*g*_. Then, the expected response to direct selection is *R*_*β*_ = *β*_*g*_ *× diag*(**G**), where *diag*(**G**) is the vector of additive genetic variance of gene expression.

We obtained five estimates of *s*_*g*_ and *β*_*g*_ for each gene and **G** among genes, one for each subdivision of the dataset. Each estimate and their significance are evaluated from the pooled posterior distributions of the three independent runs of BSFG. The estimate’s value is the mode of the joint posterior and significance is reached if the 95% HDI does not overlap with zero. We report the average mode of *s*_*g*_ and *β*_*g*_ over the five subdivisions for genes with at least one significant value in HD. We used the R package *MCMCglmm* (Hadfield, 2010) to compute the posterior modes, highest density intervals (HDI).

When comparing group of genes and their exposure to genetic selection, we used the log10 of the absolute value of *s*_*g*_ estimates.

### Weighted gene co-expression network analysis (WGCNA)

To get more insights into the genetic architecture of transcriptomes, we applied a weighted gene co-expression network analysis (WGCNA) as implemented in the R package *WGCNA* (Langfelder and Horvath, 2008). We constructed gene co-expression networks for CT and HD conditions separately at generation 1 and 22. At generation 1, we used the same data set as for the Bayesian sparse factor analysis (see main text). We applied the same general approach to data sets at G1 and G22. Prior to analysis, lowly expressed genes (minimum of 1 cpm in at least two samples) were removed. We then normalized read counts (cpm, TMM-normalized) and corrected for families and sequencing runs using the *removeBatchEffect* function in the limma R package (Pritchard et al., 2017) for each condition separately resulting in slightly different gene numbers in CT and HD at generation 1 (CT: 11,812 genes, HD: 11,890 genes). Briefly, WGCNA constructs networks using the absolute value of the Pearson’s correlation coefficient as the gene co-expression measure, which is raised to a power *β* to create the adjacency matrix. The topological overlap distance calculated from the adjacency matrix is then clustered with the average linkage hierarchical clustering. For choosing power *β* we followed the recommendations of (Zhang and Horvath, 2005) and used the scale-free topology criterion resulting in *β* = 4 in Control, *β* = 5 in Hot-Dry at G1. We used the *blockwiseModule* function in the R-package WGCNA to construct a signed network using a minimum module size of 20 and a *mergeCutHeight* of 0.1 (threshold for merging modules with highly correlated module eigengenes). Other parameters remained at their default. To test whether a module was related to offspring number, the correlation between a module eigengene (corresponds to the first principal component) and offspring number was used. Module preservation statistics were computed using the *modulePreservation* function (1000 permutations) implemented in WGCNA (Langfelder et al., 2011). Network module preservation statistics quantify how density and connectivity patterns of modules defined in a reference data set are preserved in a test data set without the need to define modules in the test data set. The composite measure *Zsummary* and *medianRank* were used to evaluate module preservation in other conditions. *Zsummary* is a composite statistic that combines preservation of density as well as connectivity of a module. Since *Zsummary* strongly depends on module size, it is only appropriate to examine preservation of the same module in different conditions. Values below two indicate no preservation, values between two and ten moderate and above ten strong preservation. To compare different modules within the same condition *medianRank* (Figure S1) is more informative which gives relative preservation of modules (Langfelder et al., 2011). For hub gene identification we used the *intramodularConnectivity* function of the WGCNA R package and selected the top ten genes of each model showing the highest intramodular connectivity. While the choice of ten was arbitrary, we tested different cutoffs, and the overall results were consistent. We then tested whether these hub genes were enriched in eQTLs, and whether they significantly differed from other genes in selection and evolutionary responses.

### RNA-seq single nucleotide polymorphism detection

To uncover DNA sequence variation (single nucleotide polymorphisms, SNP), we developed bio-informatics pipelines (see Figure S9) to analyze our whole transcriptome RNA-seq samples. Recent studies demonstrated that SNP detection on RNA-seq samples can be accurate when following good practices and a correct set of filtering criteria. For example, Adetunji et al. (2019) and Jehl et al. (2021) have shown that 65% and 91% of SNPs found with DNA-seq in expressed regions could be detected with RNA-seq, respectively. We used GATK-4.2.3.0 (Van der Auwera and O’Connor, 2020) to detect genomic variants from the total set of 822 whole transcriptome RNA-seq samples collected from generations 1 and 22. Mapped RNA-sequences (BAM files) were edited to add read groups (samtools 1.14; Danecek et al. 2021), and MAPQ values equal to 255 were re-calibrated to 60 to match GATK format. Duplicated reads were marked (picard Duplicates function; Broad 2019), and N-cigar reads were split using GATK SplitNCigarReads function. Variant detection was done for each sample with GATK HaplotypeCaller using the parameters `-ERC GVCF –pcr-indel-model NONE –dont-use-soft-clipped-bases`. The joint variant call was performed using GATK GenomicsDBImport and GenotypesGVCFs functions with parameter –max-genotype-count 1500. Bi-allelic single nucleotide polymorphisms (SNPs) were then selected using GATK SelectVariants with parameters –select-type-to-include SNP –restrict-alleles-to BIALLELIC –exclude-non-variants. Missing genotypes (DP=0) were labeled as missing (“./.”) using bcftools-1.14 (Danecek et al., 2021), and SNP tags were re-calculated using bcftools +fill-tags function. We then removed low-quality SNPs using GATK VariantFiltration function with six filters: QD < 2, FS > 30, SOR < 40, MQ < 40, MQRankSum < −12.5 and ReadPosRankSum < −8.0. SNPs were annotated on the reference genome using snpEff-5.1 software (Cingolani et al. 2012; default interval size of 5kb) resulting in 566,296 quality-checked SNPs (Data S2). Finally, we filtered out SNPs with a call rate lower than 50% using GATK VariantFiltration function, leading to a VCF file containing 358,142 SNPs (online supporting file Data S3). The SNPs were uniformly distributed on the genome (Figure S10) and among 11,216 genes with 31.76 SNPs per gene on average (Figure S11). The correlation between average read count and number of SNPs per gene was low (Figure S12), suggesting that SNP detection was not affected by gene expression levels.

### Imputation of missing genotypes

Around 38% of the genotypes are missing in the dataset, compromising non-missing markers’ overlap between generations in each line. To impute missing genotypes, we used Beagle-5.4 software (Browning et al., 2021) with the VCF file as a sole input. Imputed markers were then merged with the original VCF file and SNP tags were re-calculated using bcftools +*fill-tags* function (files available online in Data S4). Importantly, we tested Beagle-5.4 accuracy by building a toy dataset from all SNPs with a call rate of 100% (7,844 SNPs), and by generating missing markers using the same distribution as in the complete VCF file. In all cases, more than 98% of genotypes were accurately imputed (see Supplementary Material). Imputed data available online as Data S5.

### eQTL associations

We used the imputed VCF file to calculate eQTL associations from the half-sib/full-sib cross at generation 1 in HD. SNPs with a minor allele frequency (MAF) lower than 0.05 were filtered out, yielding a set of 141,387 SNPs (see Data S6). After applying a quantile normalization on the expression of each gene (reference: standard normal distribution), genome-wide association analyses (GWAA) were performed on a gene-wise basis to detect eQTLs for each gene expression level as a phenotype by using a linear mixed model implemented in the software GEMMA-0.98.5 (option -lmm; Zhou and Stephens 2012). GEMMA was run with the SNP-based kinship matrix (genomic relatedness matrix) as random factor. The kinship matrix was calculated with GEMMA prior to running the association analysis. As GEMMA does not accept VCF files in input, the VCF file was converted into a.bed file with PLINK2 (Chang et al., 2015). Phenotypes (gene expression data) were further added to a.fam file (PLINK2) containing the pedigree structure of the data. For each GWAA, a genomic control (GC) correction was applied to correct for p-value inflation (Bacanu et al., 2000), and an FDR of 5% was applied to detect and collect significant eQTL associations. We found 958 eQTLs significantly associated with the expression level of 556 distinct other genes. All the eQTLs were distant, *i.e*., associated with the expression level of other genes rather than their carrier gene. The same analysis was run by using relative fitness measurements as a phenotype, no significant associations were found. Results available online as Data S7.

### Detection of significant allele frequency changes and parallelism

Selection at the sequence level can be inferred from parallelism of allele frequency changes (AFCs) among replicates (Yeaman et al., 2018; Barghi et al., 2019, 2020). We used the imputed VCF file to perform the analysis of AFCs within lines between generations 1 and 22, in CT and HD separately. In CT and HD, four lines (L1, L3, L5, L6) were sequenced at generations 1 and 22. Additional lines were sequenced at G22 only: L4, Mx1, and Mx3 in CT; L2, Mx1, and Mx2 in HD. To calculate AFCs, we used allele frequencies at generation 1 when available (L1, L3, L5, L6), or we used a common pool at generation 1 (all available lines merged in a single pseudo-population). We also ensured that the addition of mixed lines was not introducing a bias in parallelism estimates used in this work. To do so we re-calculated all the statistical tests without these lines and found no significant difference (see Supplementary Material). To do so, we split the imputed VCF file into one file per line and generation for each environment, as well as two additional files with all samples at generation 1 for CT and HD. SNP tags were then re-calculated using bcftools +fill-tags function. Allele frequencies were polarized by MAF at generation 1 to match the eQTL analysis. For each line and each SNP, the allelic frequency change (AFC) was then the difference between allele frequencies at generation 22 and 1. The resulting dataset is available online as Data S8.

To detect significant AFCs in each line of HD environment, we used a distribution merging all the AFCs measured in CT environment (*i.e*., from all the pooled lines) as a single reference distribution for a non-parametric bootstrap procedure. We tested whether the tail probability of each SNP’s AFC within each HD line at G22 was larger than 97.5% based on its confidence interval derived from a sample of 10,000 bootstrapped AFC distributions from the reference AFC distribution in CT. The number of AFCs randomly sampled in each bootstrap replicate equated the number of SNPs within each HD line. This allowed us to test for significant AFC in all seven HD lines sequenced in G22. We defined AFC parallelism as the number of lines in which a SNP was showing a significant AFC. Parallelism thus varies between 0 and 7. The full results are available online as Data S9. We then defined the parallelism of a gene as the largest parallelism detected among its SNPs. We used a gene-based definition of AFC parallelism over an LD-based approach because our estimates of selection are at the gene level and analysis of LD per chromosome shows that LD-tagging would lead to similar results given the LD decay rate observed (see Figure S13).

### Functional enrichment

Three subsets of genes 1) Hub genes, 2) eQTL carriers, 3) overlap between hub genes and eQTL carriers, were tested for functional enrichment using g:Profiler version e111 eg58 p18 30541362 (Kolberg et al., 2023). We further used STRING, version 12.0 (Szklarczyk et al., 2019) to test for functional enrichment in proteins after converting genes to proteins using g:profiler. The results of the enrichment analysis can be found online as Data S10.

## Results

### Evolutionary changes are consistent with predictions

We have estimated the expected evolutionary change in mean expression level of each expressed gene as *s*_*g*_ = *cov*_*A*_(*z, w*), the genetic covariance of the phenotype *z* (here gene expression) and relative fitness *w* (Robertson, 1966; Price, 1970), hereafter called *genetic selection*. Since evolutionary changes require heritable genetic variation, we have also estimated the additive genetic variance of expression of each transcript (*V*_*A*_). Genes greatly varied in their proportion of heritable genetic (*V*_*A*_) to phenotypic variance (*V*_*P*_) as shown by the distribution of their heritability coefficients 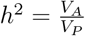 (Figure S2). Median *h*^2^ in HD was 0.64, very close but significantly lower than in CT (0.66, *P* < 0.0001, permutation test), indicating a larger genetic contribution to phenotypic variation in expression levels than environmental noise.

The estimates of *s*_*g*_ in HD were predictive of the observed evolutionary changes in gene expression in HD: genes under genetic selection to increase expression in generation 1 (G1) increased in gene expression in generation 22 (G22) and those under selection for reduced expression decreased in gene expression. The average expression change of positively selected genes was significantly positive (0.067 logFC; bootstrap 95% CI: [0.036, 0.102]), and similarly for negatively selected genes (−0.094 logFC; bootstrap 95% CI [-0.133, −0.053]; Figure 2A). More generally, the transcriptome-wide correlation between *s*_*g*_ in HD and the observed evolutionary changes in HD was positive and significant (*ρ* = 0.22; permutation test: *P* < 0.0001). We found 55 genes with significant *s*_*g*_ in HD at generation 1 (see Methods). This set of 55 genes had an even stronger correlation between predicted and observed evolutionary changes in expression (*N* = 55: *ρ* = 0.64, permutation test: *P* < 0.0001). Based on this substantial and significant correlation we may conclude that predictability of evolutionary changes is thus high in the HD environment (Figure 2B).

**Figure 2:**
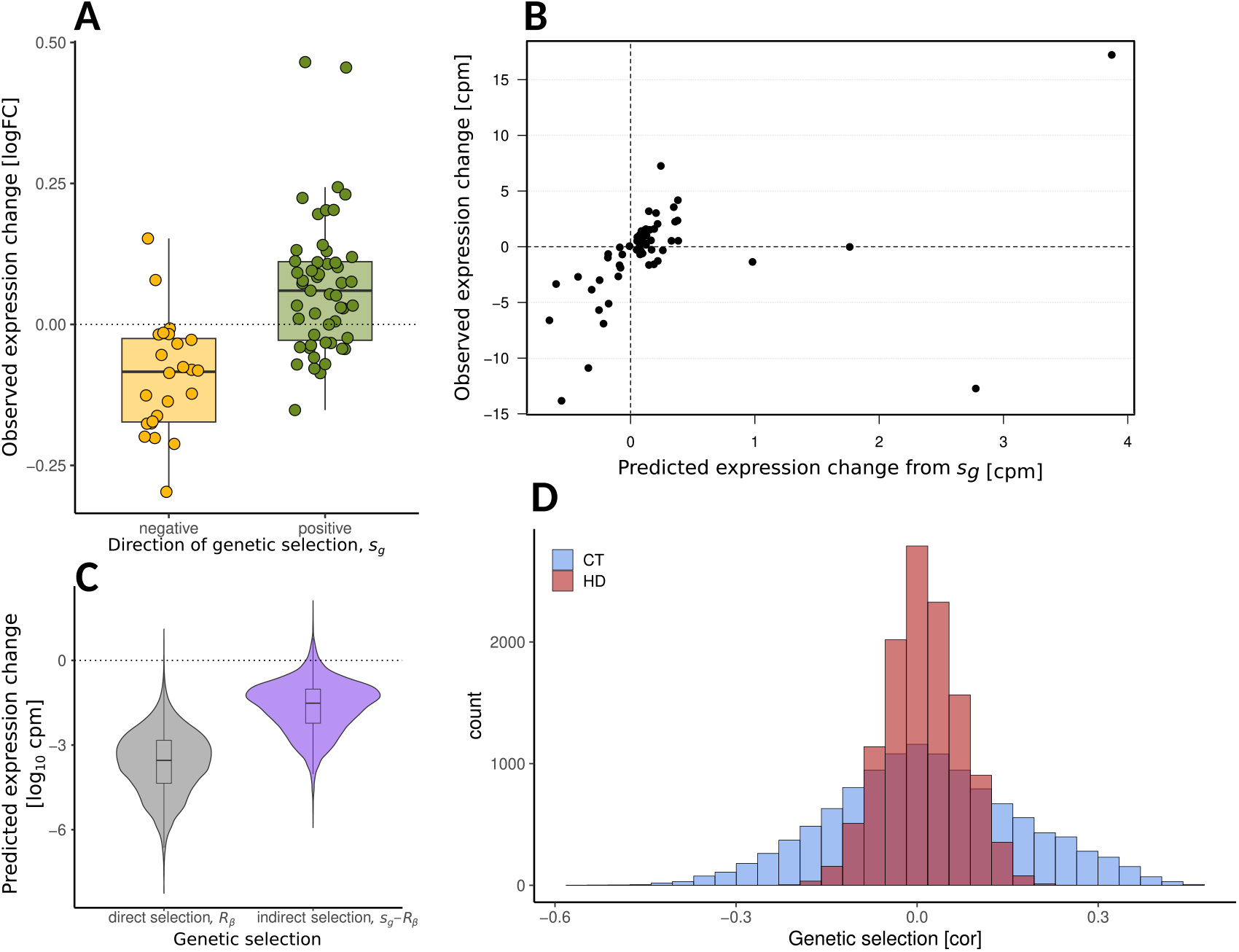
Genetic selection and evolutionary changes in HD. (**A**) Evolutionary response of gene expression (log_2_ fold change—logFC) in Hot-Dry conditions of the genes having significant genetic correlation with fitness in the first generation (genetic selection). The evolutionary response is the difference between selection lines after 20 generations of adaptation and non-adapted control lines in Hot-Dry conditions. (**B**) Relationship between predicted responses to genetic selection based on significant estimates of *s*_*g*_ and observed evolutionary changes after 20 generations of selection in Hot-Dry conditions(in counts per million gene expression levels—cpm). (**C**) Comparison of the absolute responses to direct (*R*_*β*_) and indirect (*s*_*g*_ *− R*_*β*_) selection in Hot-Dry conditions at the first generation for all 11,890 genes with expression data. (**D**) Distribution of the variance-standardized genetic selection coefficients (genetic correlation *r*_*g*_) in the Control (CT) and Hot-Dry (HD) environments at generation 1.

### Predicted evolutionary response is mainly caused by indirect selection

From our data we could estimate the predicted response of gene expression to direct selection *R*_*β*_ (see Methods) and thus evaluate its importance relative to indirect selection by comparing *R*_*β*_ to the total response to selection *s*_*g*_, which includes both direct and indirect responses. We found largely weaker estimates of the response to direct selection *R*_*β*_ than to indirect selection (*s*_*g*_ *− R*_*β*_; Figure 2C) with median ratio *R*_*β*_*/s*_*g*_ of about 0.6%, which means that indirect selection contributes most to total selection. Direct and indirect selection can be of opposite signs and may cancel each other, or lead to opposite signs between direct (*R*_*β*_) and total (*s*_*g*_) selection. In genes with *R*_*β*_ and *s*_*g*_ of same sign (75% of total) or opposite sign (25%), *R*_*β*_ contributed a similar median proportion of about 1% of *s*_*g*_ (1.02% and 1.01% respectively). Only 86 genes (0.7%) showed a larger predicted absolute response to direct selection *R*_*β*_ than to predicted total selection *s*_*g*_, of which 46 genes had opposing direct and total selection. This suggests that the response to direct selection is constrained by opposing indirect selection in only rare instances. More generally, indirect selection is in the same direction as direct selection and constitutes the majority of genetic selection *s*_*g*_. The constraining effect of indirect selection on expression evolution is thus likely small.

### Gene co-expression networks and selection

The prevalent role of indirect selection underscores the importance of genetic correlations between expression levels. Analysis of gene co-expression networks is a powerful way to assess the importance of correlations among gene responses and identify clusters of co-regulated genes. A weighted gene co-expression network analysis (WGCNA; Langfelder and Horvath 2008) on gene expression levels in HD uncovered 26 modules of co-expressed genes (Figure 3A) and nine co-expression modules in CT. Overall, CT modules were preserved in HD, *i*.*e*., there were no fundamental differences in the way gene expression levels are correlated with each other (Figure 3D). However, it seems that correlations between expression levels became weaker in HD leading to a splitting of the CT modules. We tested each module’s fitness association by correlating the module’s eigengene (corresponding to the first principal component of each module) with fitness. Six CT modules were significantly associated with fitness variation, but none of the HD modules were (Figure 3C). Additionally, when comparing selection coefficients of single gene expression levels between CT and HD, we observed a decrease of the number of significant coefficients from 729 in CT to 55 in HD, and a narrower distribution in HD relative to CT (Figure 2D), pointing to weaker genetic selection in HD. Therefore, it seems that the stress imposed by the HD conditions caused a weakening of expression-fitness and module-fitness associations, alongside a decrease of mean fitness in HD (from 49.1 in CT to 25.7 surviving offspring per female; see Koch and Guillaume, 2020a).

**Figure 3:**
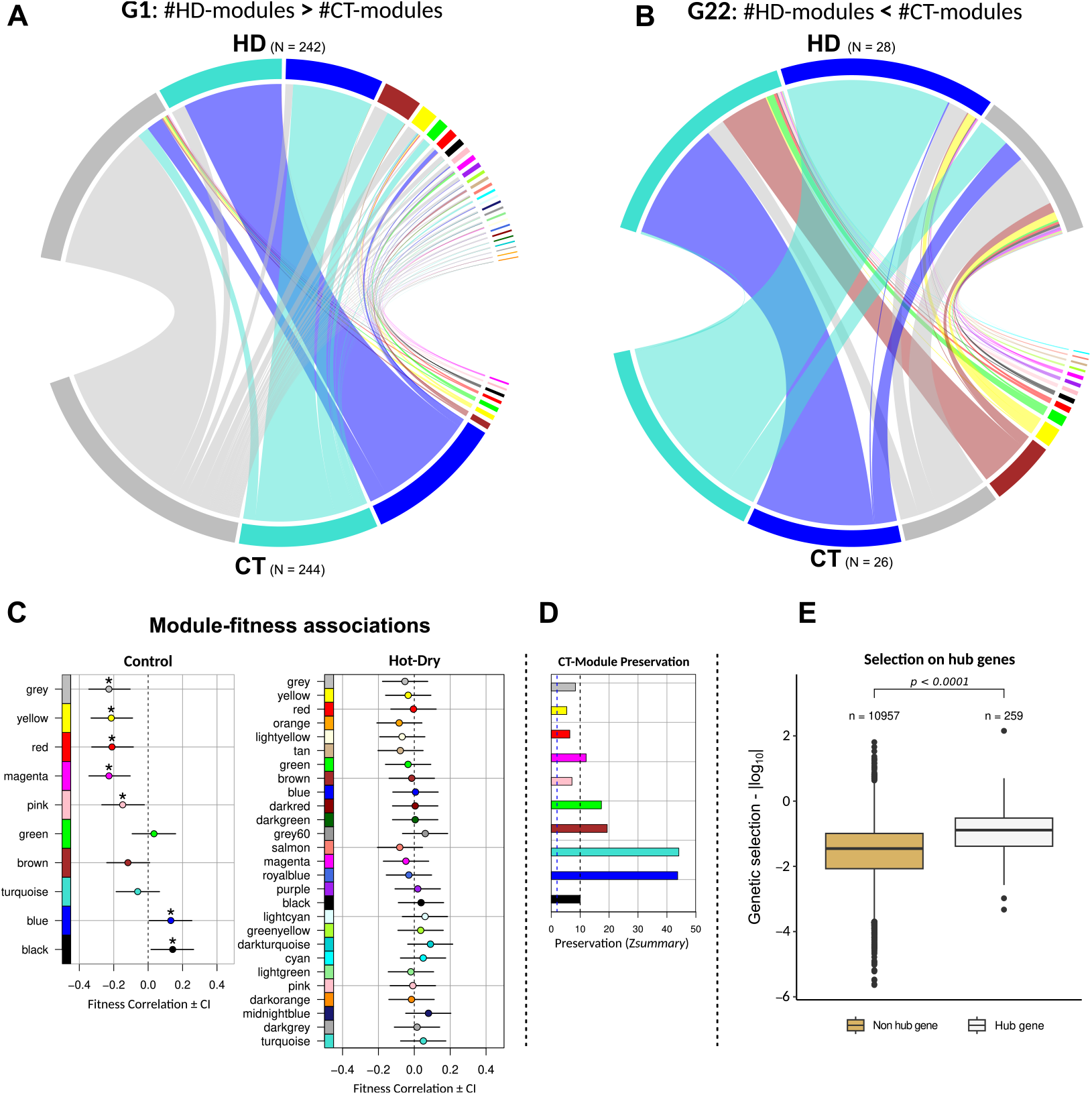
Gene co-expression network evolution in CT and HD. (**A**) at generation 1 and (**B**) generation 22. The number of sequenced individuals for each analysis is shown on the graphs. Colours refer to gene co-expression modules within each environment (same colours in different conditions refer to independent modules). Assignment of colours to modules is based on their size, *e.g*., the largest module is always ‘turquoise’. The modules’ overlap and gene number are presented in Figures S3 (G1) and S4 (G22). The grey modules correspond to genes not assigned to any cluster of co-expressed genes. Expression levels of genes assigned to the different modules in HD are shown in Figure S8. Unassigned genes (grey module) showed the lowest expression. (**C**) Fitness association of the gene co-expression networks in HD and CT shown in A, and (**D**) preservation of CT modules in HD at generation 1. Zsummary is a composite statistic that combines preservation of density as well as connectivity of a module. The dashed line at 2.0 indicates the thresholds between no and weak preservation; the line at 10 gives the threshold between moderate and strong preservation. (**E**) Difference in absolute log(*s*_*g*_) of the hub versus non-hub genes in HD. A comparison of expression levels between hub and non-hub genes is shown in Figure S7.

Reassessing the structure of gene co-expression networks after 20 generations of adaptation, we found that the number of modules in HD was drastically reduced to two modules in the adapted lines while the number of CT modules in the CT lines remained high at 14 (Figure 3B). We thus found the reverse pattern of integration of sub-networks into two large modules at generation 22 in HD, also kept for HD lines in CT (Figure S6). Note that the analyses were done on 28 (26) samples in HD (CT). However, downsampling the WGCNA in generation 1 to similar sample sizes led to higher numbers of modules in HD, not lower (on average 55 modules with 50 replicates). Moreover, the previous network fragmentation caused by exposing CT lines to HD conditions was somewhat recovered at generation 22 (module number increased from 14 to 19 for CT lines transplanted to HD; see Figure S5). Therefore, we can conclude that evolutionary changes which occurred during adaptation to HD conditions have led to a rewiring (change of topology) of gene co-expression networks.

### Network position is associated with selection and evolutionary change

Despite a weaker module association with fitness at the gene network level, we found that the 260 hub genes in HD (10 most connected genes within each module) were associated with significantly higher (4-fold increase) absolute genetic selection *s*_*g*_ compared to non-hub genes in HD (median *s*_*g*_ non-hub genes: 0.03, hub genes: 0.122; Cliff’s delta = 0.46; *P* < 10^−4^, permutation test, Figure 3E). Consistent with this observation, hub genes showed more evolutionary changes in expression compared to non-hub genes; they were over-represented among genes with significant evolutionary changes (*P* = 1.1 *×* 10^−3^, Fisher’s exact test). Overall, they had a slightly larger (1.3-fold) absolute evolutionary change (median logFC non-hub genes: 0.09, hub genes: 0.11; Cliff’s delta = 0.11; *P* = 1.7 *×* 10^−3^, permutation test) in generation 22. However, absolute direct selection on hub genes in the first generation was significantly weaker compared to other genes (median non-hub: 2.13e-05, hub genes: 1.01e-05; *P* < 10^−4^, permutation test). To avoid confounding effects caused by differences in expression levels between hub and non-hub genes, as well as unequal sample sizes (hub = 260 vs. non-hub = 10,957), and to confirm that differences in *s*_*g*_ and evolutionary changes are due to network position, we applied a resampling approach. Random subsets of non-hub genes matching both the number of hub genes and their expression level distribution were created. Specifically, non-hub genes were sampled to equally cover the expression level quantiles of hub genes. We then compared the distribution of 10,000 permutations of *s*_*g*_ and evolutionary changes in these sets of non-hub genes to the observed values for hub genes, finding that the observed hub gene values were outside the distribution of resampled non-hub genes (*s*_*g*_: *P* = 0.038; evolutionary changes: *P* < 10^−4^). Additionally, as a complementary approach, we calculated median *s*_*g*_ and evolutionary changes after correcting for expression levels by taking the residuals from a regression of log-transformed absolute *s*_*g*_ and log-transformed absolute evolutionary changes on expression levels. This resulted in median *s*_*g*_ values of 0.094 for non-hub and 0.118 for hub genes, and median evolutionary changes of 0.066 and 0.296, respectively. These results confirm that network position itself, independent of expression levels, is associated with *s*_*g*_ and evolutionary changes. Lastly, as additional support to the evolutionary importance of network position, we found a strong negative relationship between a gene’s genetic autonomy and its network centrality in HD’s coexpression network (Spearman *ρ* = −0.60; *p*-value < 2.2 *×* 10^−16^). Autonomy is inversely related to the square of the multiple correlation coefficient 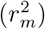 of a gene with other genes computed from our estimates of the G-matrix (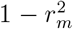; Hansen and Houle, 2008) and centrality is its network connectivity measured from phenotypic correlations (Langfelder and Horvath, 2008). This shows that more central genes are more genetically correlated to other genes and demonstrates that phenotypic correlations have a strong genetic basis.

### Selection on gene expression is accompanied by parallel allele frequency changes

To further understand the link between evolutionary changes in gene expression levels and those at the sequence level, we tested for associations between selection on gene expression (*s*_*g*_) and parallelism of allele frequency changes (AFCs) at the respective genes among replicated lines in HD. Genome-wide, 55,063 SNPs showed significant AFC in HD, when tested against the reference AFC distribution in CT (see Methods). We then defined the parallelism of a gene as the largest parallelism detected among its SNPs. This resulted in 8,906 genes being classified as exhibiting AFC in at least one line. Among those, 147 genes carried 179 highly parallel SNPs with significant AFC in at least five lines. Only one SNP had significant AFC in all seven lines sampled in HD (see Figure S14A). The correlation between *s*_*g*_ and gene parallelism was positive and significant (*ρ* = 0.215, *adj. R*^2^ = 0.046, *p*-value < 2.2e-16). Genes binned into different categories of AFC (“no shift”; “single shift”; “partially or fully parallel shifts”, Figure 4A) significantly differed in *s*_*g*_ (all two-sided Mann-Whitney-U tests *P* < 1 *×* 10^−4^; median absolute log10 *s*_*g*_: −2.02 (no shift), −1.48 (single), −1.33 (partial/full)). This corresponds to a 4.98 fold increase in absolute *s*_*g*_ between non-shifting and parallel genes, but a moderate Cliff’s delta (0.36) indicates substantial overlap between their distributions. Genes with significant genetic correlation with relative fitness (*r*_*g*_, variance-standardized *s*_*g*_) were significantly enriched in highly parallel SNPs (AFC in ≥ 5 lines; *P* = 5 *×* 10^−3^, permutation test on 2,245 SNPs in 75 genes minus five genes not carrying any SNP). These results suggest that selection on gene expression levels may play a key role in driving parallelism of AFCs among replicates.

**Figure 4:**
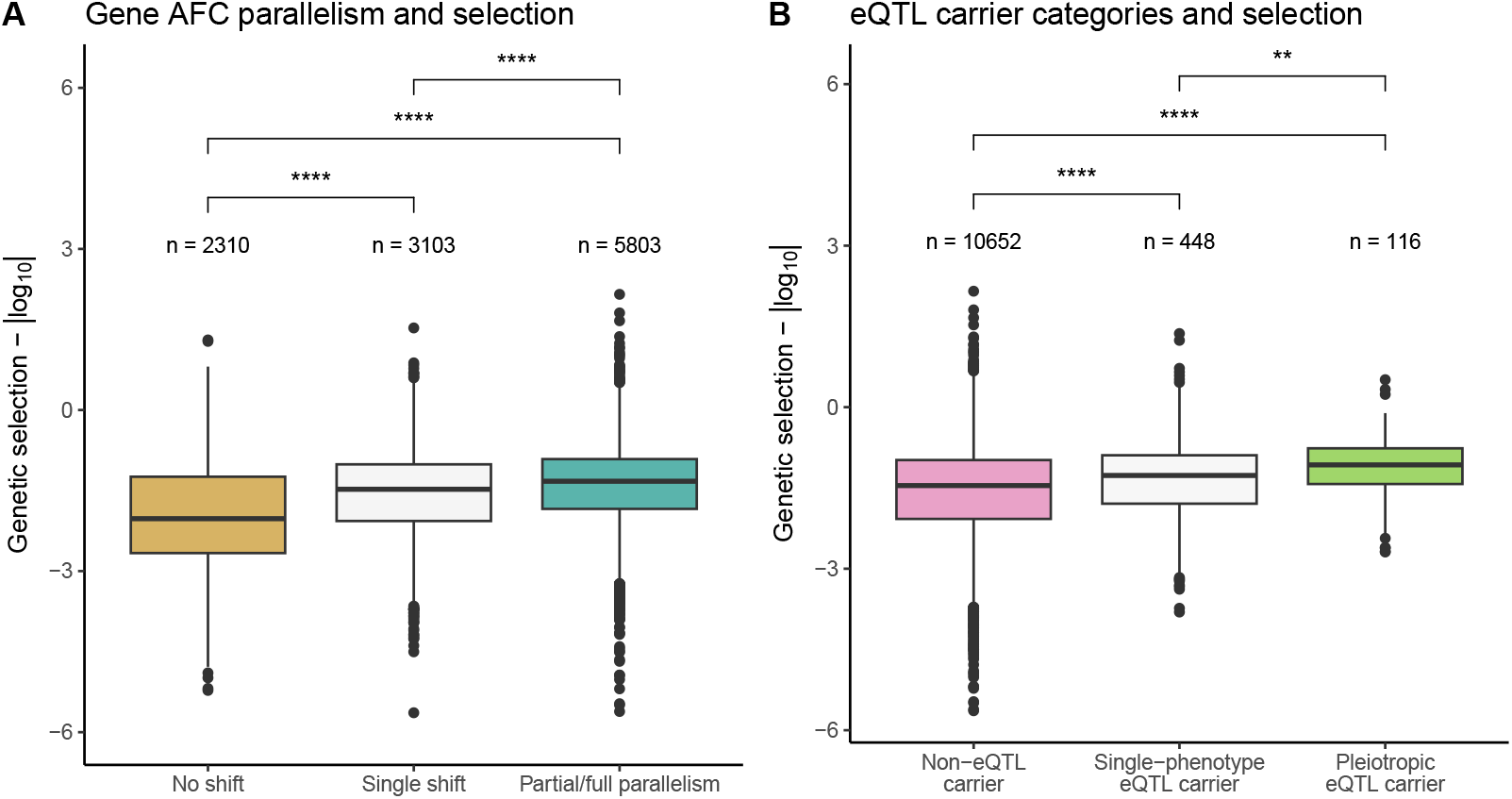
Genome-wide, gene AFC parallelism and eQTL carrier pleiotropy are significantly associated to selection at the gene expression level. For each panel, the distribution of genetic selection coefficients *s*_*g*_ (*i.e*., covariances with fitness) is plotted by category in absolute log-scale, with all sample sizes and pairwise Mann-Whitney-U tests being shown (****: P < 0.0001, **: P < 0.01). (**A**) Gene parallelism categories: “No shift”, “Single shift” and “Partial/Full parallelism”. (**B**) eQTL carrier genes categories: “non-eQTL carrier”, “non-pleiotropic eQTL carrier” (*i.e*., a single phenotype association) and “pleiotropic eQTL carrier” (≥ 2, max = 41; see Figure S15).

On the other hand, we did not find enrichment of parallel AFCs in the genes with either largest or significant evolutionary changes in gene expression; we found no enrichment in parallelism among 2,424 SNPs located in the first 100 genes with largest evolutionary response (*P* = 0.71, permutation test), nor among 2,250 SNPs belonging to the 53 genes with a significant evolutionary response (from the 55 genes found in Koch and Guillaume 2020b and carrying at least one SNP; *P* = 0.68, permutation test). This suggests that although selection on expression and parallelism of AFCs appear linked, the SNPs with parallel AFC did not seem to affect the evolution of expression of the genes to which they belong.

### Stronger signals of selection in hub genes and more pleiotropic eQTL carrier genes

Since in our case selection is mostly indirect, we expect to find stronger signals of selection within more connected genes, *i*.*e*., within hub genes or genes more functionally connected to other genes. Hub genes and genes carrying eQTLs (*i*.*e*., SNPs significantly associated to expression variation at other genes) are obvious candidates to also test for the effects of indirect selection on gene-level AFC parallelism. We found that the 259 hub genes in HD (ten per module; minus one gene not carrying any SNP) were significantly more parallel than the rest of the genome (*P* = 1.9 *×* 10^−4^, permutation test). We note that all detected eQTLs were distant, *i*.*e*., associated with the expression level of other genes rather than their carrier gene (see Material and Methods).

Among these 259 hub genes, genes with partial (AFC in two to four lines) or full parallelism (AFC in ≥ 5 lines) were over-represented compared to genes with no significant shift (*χ*^2^ = 21.34, df = 2, *p*-value = 2.323e-05). Additionally, we found that hub genes were more likely to carry an eQTL than expected by chance (*χ*^2^ = 30.688, df = 1, *p*-value = 3.031e-08). This strengthens the vision that hub genes may be functionally linked with other expressed genes.

We then focused on the 564 genes carrying at least one eQTL. We found that eQTL carrier genes were more highly parallel (in ≥ 5 lines; *χ*^2^ = 11.974, df = 1, *p*-value = 5.396e-04). Defining the pleiotropy of one eQTL as the number of distinct phenotypes (*i*.*e*., gene expression levels) associated to it, we then asked whether more pleiotropic eQTL carrier genes were under stronger genetic selection. Using three pleiotropic categories, “non-eQTL carrier”, “non-pleiotropic eQTL carrier” (*i*.*e*., a single phenotype association) and “pleiotropic eQTL carrier” (≥ 2 associated phenotypes, max = 41; see Figure S15), we first found that expression of eQTL carrier genes was under significantly stronger genetic selection compared to non-eQTL carriers (Figure 4B; all two-sided Mann-Whitney-U tests *P* < 1 *×* 10^−4^) with a median log10 absolute *s*_*g*_ of −1.46 in non-eQTL compared to −1.23 for eQTL carriers, which corresponds to a 1.7-fold increase in absolute *s*_*g*_ from non-to eQTL carriers (Cliff’s Delta = −0.18). Second, genetic selection *s*_*g*_ was significantly higher among pleiotropic than non-pleiotropic eQTL carrier genes (Figure 4B; two-sided Mann-Whitney-U test *P* < 1 *×* 10^−2^). Median log10 absolute *s*_*g*_ increased from −1.27 for single-phenotype eQTLs to −1.07 for pleiotropic eQTLs, corresponding to a 1.6-fold increase in absolute median *s*_*g*_ (Cliff’s delta = −0.18). Although the small effect sizes reflected by the Cliff’s delta values indicate substantial overlap in *s*_*g*_ between the groups, the fold-changes and significant differences point to biologically meaningful differences.

### Functional enrichment of eQTL carrier and hub genes

We performed a Gene Ontology (GO) category enrichment analysis to gain insights into the molecular functions of hub and eQTL carrier genes, with a 5% FDR. Similar analysis on sets of genes with significant evolutionary change or highest parallel AFC yielded no significant results at 5% FDR. Apart from broad functional categories like *cellular processes* or *metabolic processes*, eQTL carrier genes showed mostly enrichment in processes linked to *nuclear division* (GO: 0000280), *development* (GO: 0032502), *reproduction* (GO: 0032502) (*e.g*., *gamete generation* (GO: 0007276), *regulation of reproductive processes* (GO: 2000241) or *meiotic cell cycle processes* (GO: 1903046)), and *stress response* (GO: 0006950). Enrichment in hub genes showed a similar pattern with *developmental processes* (GO: 0032502), *gamete generation* (GO:0007276) or *sexual reproduction* (GO: 0019953) being among the most significantly enriched categories. This may underline that biological processes in females are targeted towards increasing the number of offspring. Interestingly, both groups, hubs and eQTL carrier genes, showed one of the strongest enrichment for *epoxide hydrolase activity* (GO: 0004301), which was also the top enrichment category when we tested genes acting as both, hub and eQTL carrier genes. In insects, epoxide hydrolases are mostly known for their involvement in the metabolism of juvenile hormones (Daimon and Shinoda, 2013). Given the profound role of juvenile hormones as regulators of gene expression (Nijhout, 1994) and their involvement in development, reproduction and life-history trade-offs (Flatt et al., 2005), we should expect them to be targets of selection and occupy central positions in gene regulatory networks.

## Discussion

We could provide a direct test of the predictability of selection responses of gene expression traits over multiple generations in the red flour beetle *T. castaneum* adapting to heat and drought stress. Our estimates of genetic selection predicted phenotypic responses in expression during adaptation. We also found stronger selection and larger evolutionary responses at highly connected hub genes occupying central positions in modules of co-expressed genes. This is in line with our analyses showing that genetic selection was mostly indirect, thus caused by selection acting on sets of correlated genes. Disentangling direct and indirect effects contributing to genetic selection revealed that direct selection contributed very little to predicted changes in expression. Instead, gene expression evolution can be better understood from a gene expression network perspective: Genes are evolving because they are highly interconnected and thus subject to high amounts of indirect selection.

Although the network context of gene expression evolution is generally expected (McGraw et al., 2011; Innocenti and Chenoweth, 2013; Josephs et al., 2017), the prevalent role of indirect selection has scarcely been shown previously. Indeed, our study is one among very few which actually measured indirect selection acting on gene expression traits within species. Houle and Cheng (2021) showed how genetic correlation among gene expression traits may play an important role in the evolution of sexual dimorphism in expression in *Drosophila melanogaster* and Groen et al. (2020) predicted substantial constraints on responses to selection of expression traits in rice caused by indirect selection pressures of opposite direction. In contrast to those studies, we could estimate indirect selection from genetic selection gradients instead of phenotypic gradients, which gives us better estimates of selection and thus better predictability of evolutionary responses (Morrissey et al., 2012; Stinchcombe et al., 2014).

Our genetic selection estimates were also informative of the parallelism of allele frequency changes (AFC parallelism) among experimental lines. Previous studies estimated either phenotypic selection on expression traits (selection differentials; McGuigan et al., 2014; Groen et al., 2020; Koch and Guillaume, 2020a; Ahmad et al., 2021) or AFC parallelism in sequence data from replicated populations (*e.g*., Bergland et al., 2014; Barghi et al., 2019; Rudman et al., 2022) but none confirmed that selection resulted in expression or allele frequency changes.

Joining population and quantitative genetics approaches, we could show that AFC parallelism at the SNP level is linked to genetic selection at the phenotypic level, on gene expression. However, the link is not direct since AFC parallelism is not correlated with evolutionary changes in gene expression of the SNP carriers. Importantly, though, we found that signals of selection were concentrated on hub genes, which were more likely to carry expression QTLs (eQTLs), showed the highest amount of indirect selection, and were enriched in evolutionary changes at the sequence (AFC parallelism) and phenotypic levels. In addition, the eQTLs with higher pleiotropy, *i*.*e*., influencing a larger number of expression phenotypes, were also under stronger selection. This shows how indirect selection induced evolutionary changes and confirms that selection primarily acts through indirect effects by targeting highly connected (pleiotropic) genes potentially affecting expression levels of many genes in expression modules.

### Constraints on transcriptome evolution

Previously, analyses of within population genetic variation reported purifying selection on highly connected genes (Josephs et al., 2017; Mähler et al., 2017) and predominantly stabilizing selection on gene expression variation (Josephs et al., 2015; Kita et al., 2017). Similarly among species, highly connected genes within networks were often found to show signs of constrained sequence evolution during divergence according to their pattern of genetic co-variation (Fraser et al., 2002; Hahn and Kern, 2005; Innocenti and Chenoweth, 2013). Considering that the link between connectedness in gene networks and pleiotropy is plausible (He and Zhang, 2006), these results are in line with the general expectation that genetic variation at more pleiotropic genes is more likely deleterious (Orr, 2000; Otto, 2004), and more so in populations under stabilizing selection at mutation-selection balance on multidimensional phenotypic optima (Martin and Lenormand, 2006).

In contrast, our study shows that selection can lead to larger evolutionary changes at more connected genes. This outcome is also expected from models of adaptation in multidimensional trait space but when the population is away from its phenotypic optimum (Fisher, 1930; Orr, 2000; Martin and Lenormand, 2006). Selection in our experimental lines was measured at the first generation of exposure to the environmental stress and evolutionary changes were measured after 20 generations of adaptation. Our measurements were thus made during the early, less constrained phase of adaptation where we expect adaptive substitutions to have larger and more beneficial effects than when the population is at its optimum (Orr, 2000; Martin and Lenormand, 2006). Furthermore, larger adaptive steps in multidimensional phenotypic space towards an optimum may be achieved by variants at more pleiotropic genes and thus favored early during adaptation. This explains why selection and evolutionary changes were stronger at hub genes in our experiment. It also explains in part why selection was generally more indirect than direct in our data, because affecting more pleiotropic genes during early adaptive steps.

From our results, we can conclude that more connected, and hence more pleiotropic genes are under stronger selection during adaptation to heat and drought stress in *T. castaneum* and thus display larger evolutionary changes. Similar results have been obtained in very different organisms and environments, ranging from bacteria in the laboratory (Ruelens et al., 2023) to trees adapting to environmental conditions in natural populations (Whiting et al., 2024). They point to the importance for (local) adaptation of larger effects on fitness or phenotypes that are possible at pleiotropic genes. While deleterious under stabilizing selection, those effects are beneficial during adaptation to new environments in micro-organisms (McGee et al., 2016; Maddamsetti et al., 2017; Ruelens et al., 2023) and more complex organisms (Rennison and Peichel, 2022; Thorhölludottir et al., 2023) or favored during adaptation with gene-flow in trees (Whiting et al., 2024). It thus emerges that pleiotropy and here centrality of genes in gene co-expression networks play a fundamental, positive role in the process of adaptation. Moreover, our study demonstrates that gene expression evolution is predictable, *i.e*. there is a substantial and significant correlation between predicted and observed evolutionary changes, during rapid adaptation and driven by indirect selection pressures on correlated expression phenotypes.

It is worth noting that genetic covariation was estimated from a half-sib/full-sib cross in common controlled environments, so most observed correlations likely reflect genetic rather than environmental effects. The strong concordance between phenotypic and genetic correlations supports our network-based conclusions: genes occupying central positions are expected to experience stronger indirect selection, consistent with the larger evolutionary changes observed for highly connected genes compared with those less connected.

### Network structure increases evolvability

Since gene expression traits are often organized into functional modules, exhibiting little correlation with genes outside their respective groups (Ihmels et al., 2002; Barabási and Oltvai, 2004), they are likely to experience concerted selection pressures if selection acts on the functional properties of the modules. Such alignment of selection with genetic co-variation can lead to unconstrained patterns of evolutionary response, and thus increase evolvability (Hansen and Houle, 2008; Walsh and Blows, 2009; Chevin et al., 2010). The alignment can be achieved if past selection has either sifted through existing genetic variation or selected for specific mutational correlation at pleiotropic genes (Chebib and Guillaume, 2022; Petit et al., 2023). Both phenomena would generate patterns of genetic covariation aligned with correlational selection, which then enhances selection responses and accelerates evolution in the direction of selection. This alignment is expected from theory showing that correlational selection can orient genetic covariance with the direction of selection leading to stronger responses to selection (Arnold et al., 2008; Chebib and Guillaume, 2021, 2022).

In line with that, several gene co-expression modules in the ancestral control conditions (CT) exhibited significant phenotypic correlations with fitness, indicating selection acting on whole-module expression and alignment of covariation patterns of expression with the direction of selection. We should thus expect that selection in the past has shaped the genetic covariance between expression levels to be aligned with selection acting on them (Watson et al., 2014; Petit et al., 2023). However, in the hot-dry treatment (HD), the phenotype-fitness correlation dwindled, showing a potentially strong effect of the stress environment on the network structure. Therefore, our results suggest that the metabolic and physiological stress responses in the first generation of selection affected gene expression in such a way as to mis-align gene expression modules with the direction of selection. Yet, evolution proceeded from that existing variation and led to changes in gene expression and network structure, potentially re-aligning co-expression modules with new selection pressures. Our results further suggest that evolutionary changes at hub genes of past modules were instrumental for the re-organization of the gene co-expression networks during rapid adaptation to the new environment. Our results thus also support the idea that the topology of gene networks is dynamical (Luscombe et al., 2004) and should reflect the topology of direct and indirect selection pressures (Burban et al., 2021; Petit et al., 2023).

### The genotype-phenotype-fitness map of adaptation in T. castaneum

Our integrated population and quantitative genetics approach allowed us to uncover some of the complexity of the molecular genetic and phenotypic underpinnings of adaptation to heat and drought stress of the red flour beetle *T. castaneum*. We have based our analyses on several hundreds of whole-transcriptomes mRNA-seq libraries extracted from whole-body mRNA in adult females. This large sampling allowed us to take advantage of a quantitative genetics approach which revealed a large genetic basis to the inter-individual variation in gene expression levels: The majority of the genes with detectable expression were only little influenced by non-genetic effects (*e.g*., cellular noise).

Our estimates of expression levels are thus sums of expression over cell and tissue specific expression. This may cause false negatives in differential expression analyses if changes in gene expression cancel each others across tissues (Johnson et al., 2013). Expression trade-offs may have caused lower power when detecting evolutionary changes in expression in generation 22, although small sample sizes may have affected power more. Additionally, genes whose expression constitutes a small part of whole-body expression may be more difficult to detect, especially if selection causes a reduction in expression levels (Johnson et al., 2013; Hoedjes et al., 2024). Estimates of selection are similarly affected since they depend on detectable inter-individual variation in gene expression. They are, however, not misleading about the nature of selection since genetic selection includes indirect effects brought by co-expression of genes within and among cells. Co-expression may be causal when genes are co-selected, for *e.g*., in the same pathways or non-causal when expression co-occur within a particular structure (tissue, organ) whose function is under selection. Relative to indirect selection, it does not matter whether variation in total expression is caused by variation in per-cell expression or variation in the total number of cells per structure (Hoedjes et al., 2024). If, for instance, a gene product is under positive selection, selection may act on any mechanism increasing the abundance of this product including increase in cell number. Change in organ allometries may result as a response to that selection, as for example in *Drosophila* females which evolved enlarged ovaries when adapting to heat stress (Hsu et al., 2020). A similar response may happen in *Tribolium* as suggested by our functional enrichment analysis. Genetic selection thus accounts for both causal (*e.g*., regulatory) and non-causal (*e.g*., allometry) sources of co-expression evolution. In sum, while whole-body mRNA-seq is likely to generate false negatives in the estimates of differential expression and selection, it is unlikely to cause false positives, making our approach conservative.

We based our genotype-phenotype mapping on partial genomic data from RNA-seq and have attempted to find signals of selection at the SNP and gene levels from the parallelism of SNP’s AFCs among experimental lines. Although this approach is a standard in the field (*e.g*., Orozco-terWengel et al., 2012; Yeaman et al., 2018), our experimental design differed significantly from previous studies by having maintained control lines throughout the whole experiment. This allowed us to take advantage of the distribution of AFCs in the CT lines as our null expectation and to account for AFCs driven by both genetic drift and selection caused by our experimental approach (*e.g*., rearing method), inevitably common to all experimental conditions. The advantage of our approach is that significance does not depend on an approximate significance threshold based on simulated p-values from models with unknown fit to our data (*e.g*., Yeaman et al., 2018; Barghi et al., 2019). However, similar to all existing approaches, it is bound to have high false negative rates for polygenic adaptation where small AFCs per genes or SNP are expected (Sella and Barton, 2019; Barghi et al., 2020). On the other hand, we do not expect increased false positive rates for signals of selection among replicated lines because parallelism is unlikely caused by random genetic drift. Here too our approach is conservative.

In any case, our joined quantitative and population genetics approach allowed us to show that AFC parallelism of expressed genes was positively associated with the intensity of selection on their expression levels. However, the genes showing evolutionary changes in gene expression were not associated with stronger AFC parallelism at the SNPs they carried, although expression changes were predicted by genetic selection. We could thus not link sequence with expression evolution within genes. Similarly, eQTL carrier genes were enriched in AFC parallelism although they were associated with expression at other genes (“*trans*” acting). These results indicate that selection acts on two levels of variation: variation within expressed sequences linked to expression regulation of other genes and phenotypic variation in expression. They also surely indicate presence of selection on regulatory variation outside expressed sequences that we do not capture with our RNA-seq approach. An additional explanation for a lack of change in expression levels despite allele frequency changes could be a change in gene product properties that is expression independent, as for instance a change of enzyme activity or affinity favoured by selection in the new environment (*e.g*., Brown et al., 2009). Future DNA sequencing should allow us to better understand the importance of regulatory variation as well as selection on allele specific expression levels.

In conclusion, through the integration of gene expression, fitness, and allele frequency data, we could build a genotype-phenotype-fitness map of adaptation to heat and drought stress in *T. castaneum* and gain valuable insights into the mechanisms underlying evolutionary changes during adaptation. Our findings emphasize the importance of measuring selection acting on gene expression variation and the critical information on the molecular basis of adaptation that can be gained from such measurements. Moreover, and in contrast to previous studies, we could demonstrate the evolutionary significance of our estimates of genetic selection by testing for the evolutionary responses of the genes under selection. This lends support to the general approach of using estimates of selection as a means to uncover the genetic and molecular architecture of adaptation. Our study highlights the importance of considering genetic correlations and network structures for predicting and understanding gene expression evolution and adaptation. More similar studies would confirm the generality of our findings.

## Acknowledgments

We thank Maria Domenica Moccia, Elke Karaus, Martina Berchtold, Encarnación Lozano for RNA extraction and quality control, Lucy Poveda and Maria Domenica Moccia for advice in library preparation and sequencing, Lennart Opitz for helping to process the RNA-seq data. We also thank Outi Savolainen, Tanja Pyhäjärvi, Chedly Kastally and Pierre Nouhaud for helpful suggestions and discussions, and Hannu Mäkinen (BioData Analytics unit, University of Helsinki) for his help with WGCNA analyses. Computations were performed on the Puhti cluster at the Finnish Center for Scientific Computing (CSC).

## Data accessibility

All the supplementary data and scripts, including the complete dataset for the genomics analysis, are available on Zenodo at https://doi.org/10.5281/zenodo.15298510. The complete transcriptomics pipeline and analysis scripts related to the genomics analysis are freely available on Github at https://github.com/charlesrocabert/Koch-et-al-Predictability-of-Gene-Expression. Sequence reads of the G1 samples are available on GEO (90 samples) and ENA (396 samples) platforms with accession numbers GSE133570 (https://www.ncbi.nlm.nih.gov/geo/query/acc.cgi?acc=GSE133570) and PRJEB76655 (https://www.ebi.ac.uk/ena/browser/view/PRJEB76655) respectively. Sequence reads of the G22 samples are accessible at GEO, with accession number GSE156256 (https://www.ncbi.nlm.nih.gov/geo/query/acc.cgi?acc=GSE156256). Sample accession numbers and description are provided in a supplementary data table on the Zenodo repository.

## Contributions

E.L.K. and F.G. designed research; E.L.K., C.R., C.B.R., and F.G. performed research; E.L.K., C.R., and C.B.R. analyzed data; and E.L.K., C.R., and F.G. wrote the paper.

## Supplementary Information to

## Supplementary analyses

### Imputation quality testing with Beagle-5.4 software on the raw VCF file

To test capability of Beagle-5.4 (Browning et al., 2021) to predict missing genotypes with our dataset, we used the raw VCF file obtained from the *Tribolium castaneum* transcriptomics data to evaluate the success rate of genotypes imputation.

**Supplementary analysis figure 1:**
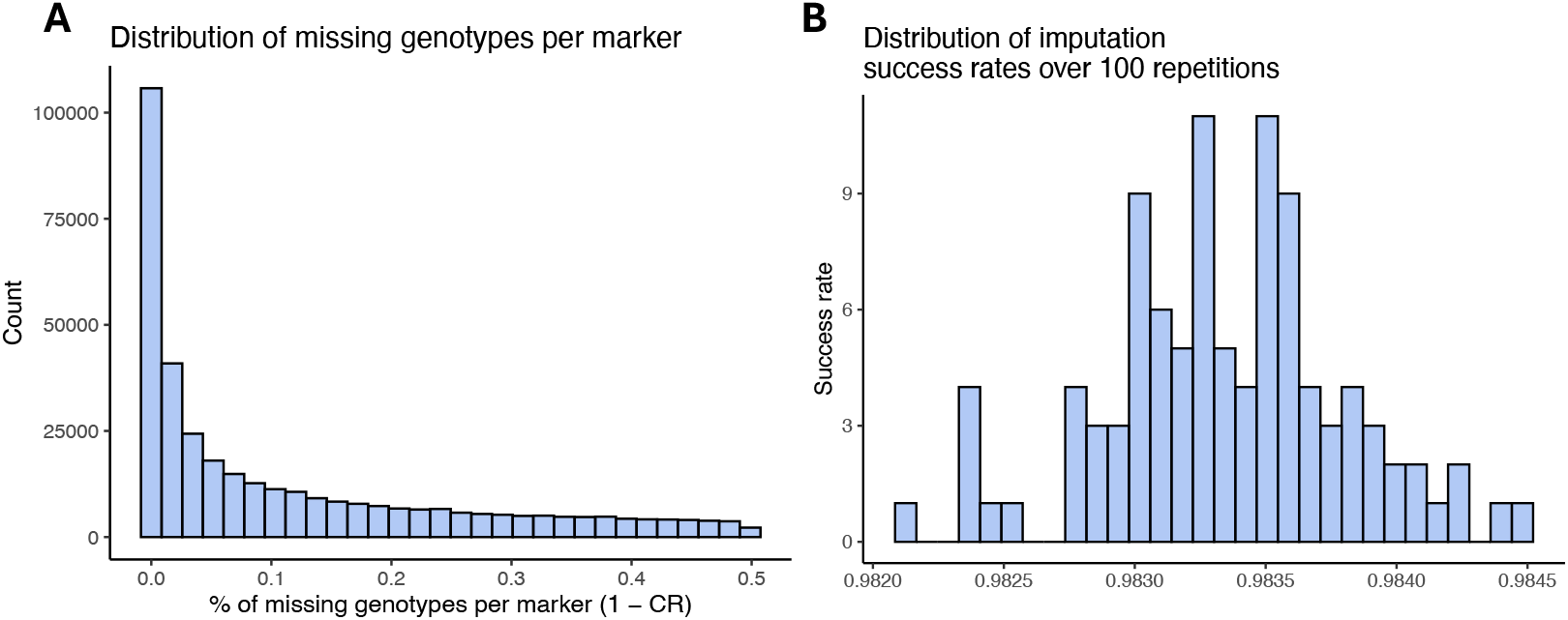
Beagle-5.4 achieves high accuracy in imputing missing genotypes in *Tribolium castaneum* transcriptomics data. (**A**) Distribution of the ratio of missing genotypes per marker, for the 358,142 markers present in the original VCF file. As SNPs have been filtered to keep markers with a call rate (CR) *>* 50% prior to this analysis, the maximum ratio of missing genotypes is 0.5. (**B**) Distribution of the genotype imputation success rate for 100 independent tests. The success rate is the ratio of missing genotypes correctly imputed (in mean, 652,600 missing genotypes are generated by artificial dataset). Mean success rate: ∼0.98% (standard error *s*.*e*. = 4.62 *×* 10^−6^).

To this aim, we selected in the raw VCF file all the markers having a call rate of 100% to create a subset of SNPs with no missing genotypes (7,837 markers out of 358,142, *i*.*e*., 2.19% of the initial VCF file). We then used this subset to generate 100 artificial datasets where missing genotypes were introduced at random following the same probability distribution per marker than in the initial VCF file (see supplementary analysis fig. 1A). For each artificial dataset, we called Beagle-5.4 with the reconstructed VCF file as a sole input. For each artificial dataset, the success rate was calculated as the ratio of imputed genotypes identical to the original ones.

We found that, in mean, the success rate was ∼0.98% (standard error *s*.*e*. = 4.62 × 10^−6^; see supplementary analysis fig. 1B). In mean, Beagle-5.14 software did not impute correctly 10,915 genotypes out of 652,600 missing genotypes.

### Testing effect of mixed lines on measures of parallelism

Mixed lines Mx1 and Mx2 were used with non-mixed lines to quantify parallelism. Since mixed lines may not be independent of non-mixed lines, we have tested whether these lines can bias our estimate of parallelism. After removing the two lines from the analysis (leading to 5 degrees of parallelism instead of 7), we found that there is no significant change in our results (as shown below).

**Supplementary analysis figure 2:**
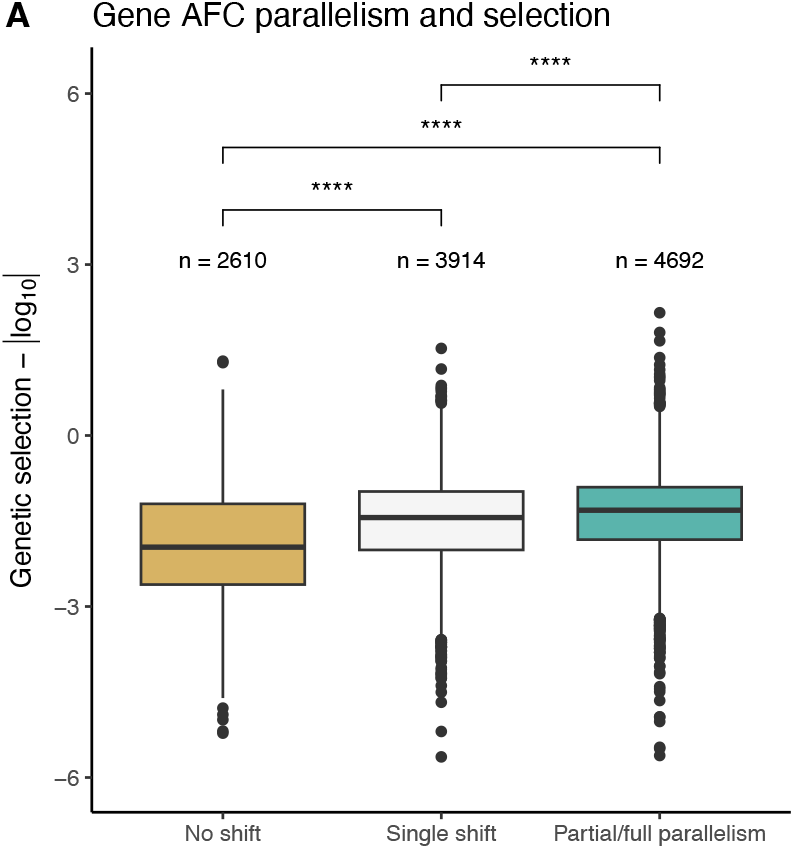
Genome-wide, gene AFC parallelism are significantly associated to selection at the gene expression level, even when removing mixed lines. (**A**) Gene parallelism categories: “No shift”, “Single shift” and “Partial/Full parallelism”. the distribution of genetic selection coefficients *s*_*g*_ (*i*.*e*., covariances with fitness) is plotted by category in absolute log-scale, with all sample sizes and pairwise Mann-Whitney-U tests being shown (****: P < 0.0001, **: P < 0.01).

Statistical tests are unchanged in main-text Figure 2 (parallelism degree vs. genetic selection, see supplementary analysis fig. 2). The 75 genes with significant genetic correlation with relative fitness *r*_*g*_ are still highly significantly enriched in highly parallel SNPs (*P* = 0, permutation test). Hub genes are still significantly more parallel than the rest of the genome (*P* = 3*×*10^−3^, permutation test), and genes with partial or full parallelism are still over-represented in this group (*χ*^2^ *p*-value = 2e-06). Finally, eQTL carrier genes are still more highly parallel (*χ*^2^ *p*-value = 2.7e-04). Thus all main results are preserved when removing mixed lines. The analysis can be re-run by setting the option “mixed lines” to FALSE in the Rscript “1 BuildDatasets.R” (see Data accessibility).

## Supplementary figures

**Supplementary figure 1:**
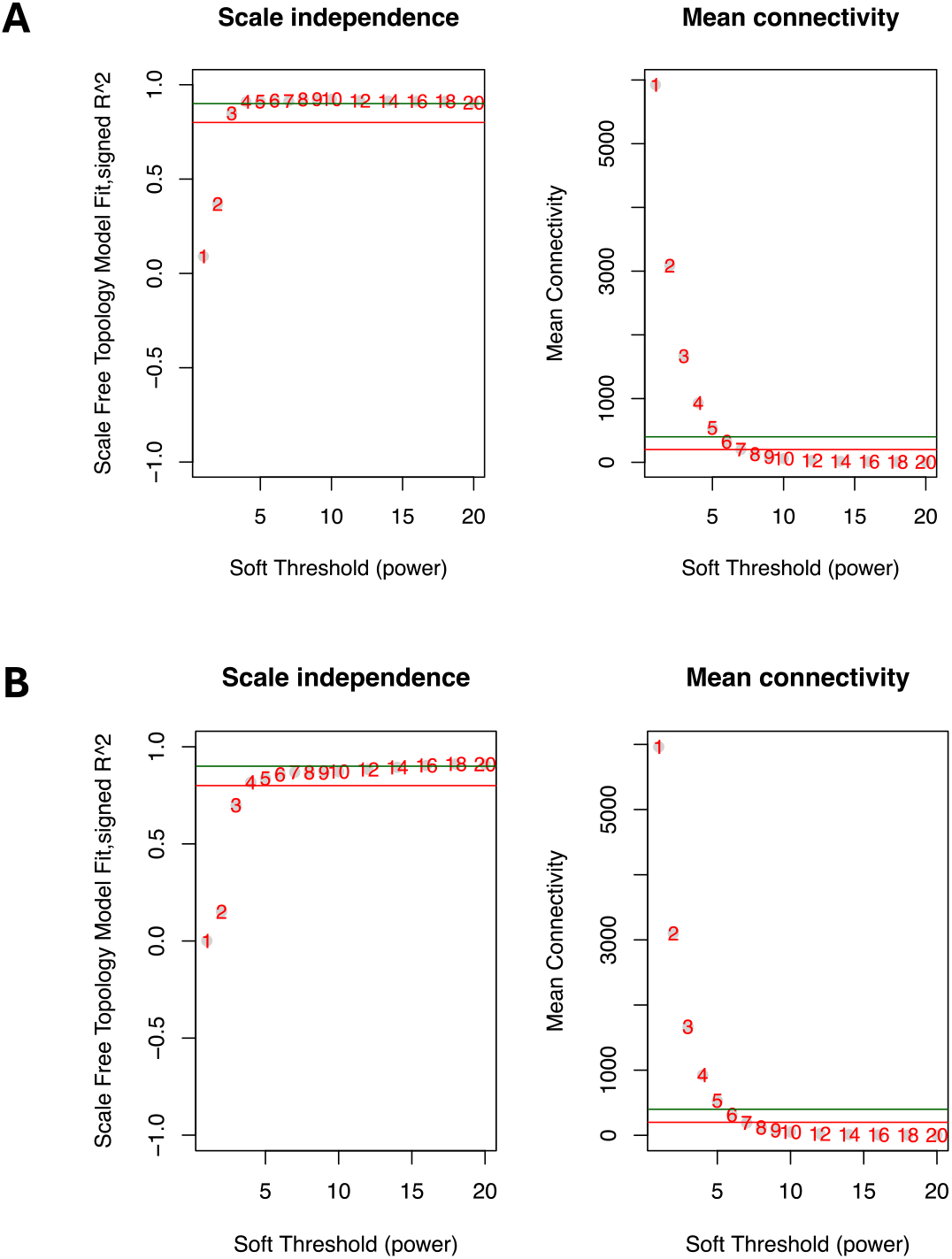
Summary network indices (y-axes) as functions of the soft thresholding power (x-axes) for Control (**A** - upper row), and Hot-Dry (**B** - lower row). Numbers in the plots indicate the corresponding soft thresholding powers. The plots indicate that approximate scale-free topology is attained around the soft-thresholding power of four for Control and for six in Hot-Dry. Red line marks *R*^2^ = 0.8, the green line *R*^2^ = 0.9 in the scale-free topology plots. Langfelder and Horvath (2008) suggested a minimum of 0.8. Because the summary connectivity measures decline steeply with increasing soft-thresholding power, it is advantageous to choose the lowest power that satisfies the approximate scale-free topology criterion. In the ‘Mean connectivity’ plots, the red line indicates 200, the green line 400.

**Supplementary figure 2:**
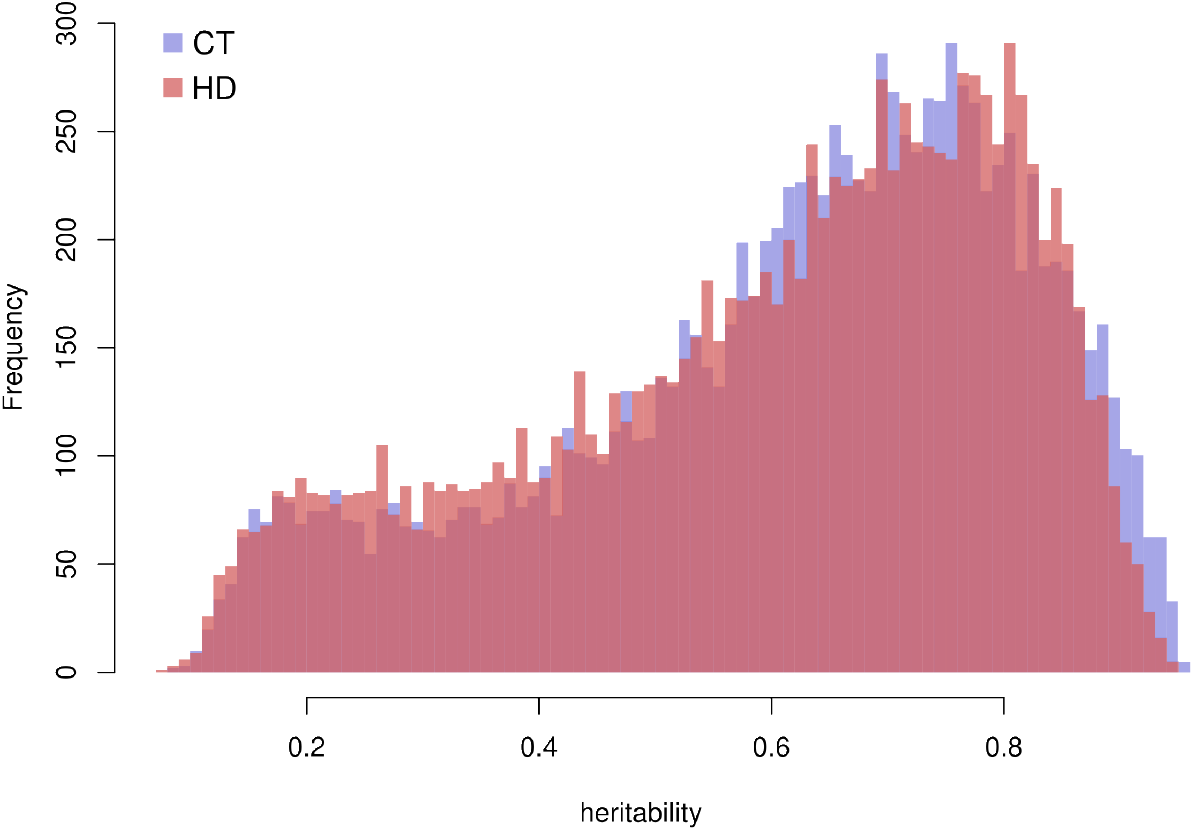
Transcriptome-wide distributions of heritability *h*^2^ of expression levels in CT (blue) and HD (red).

**Supplementary figure 3:**
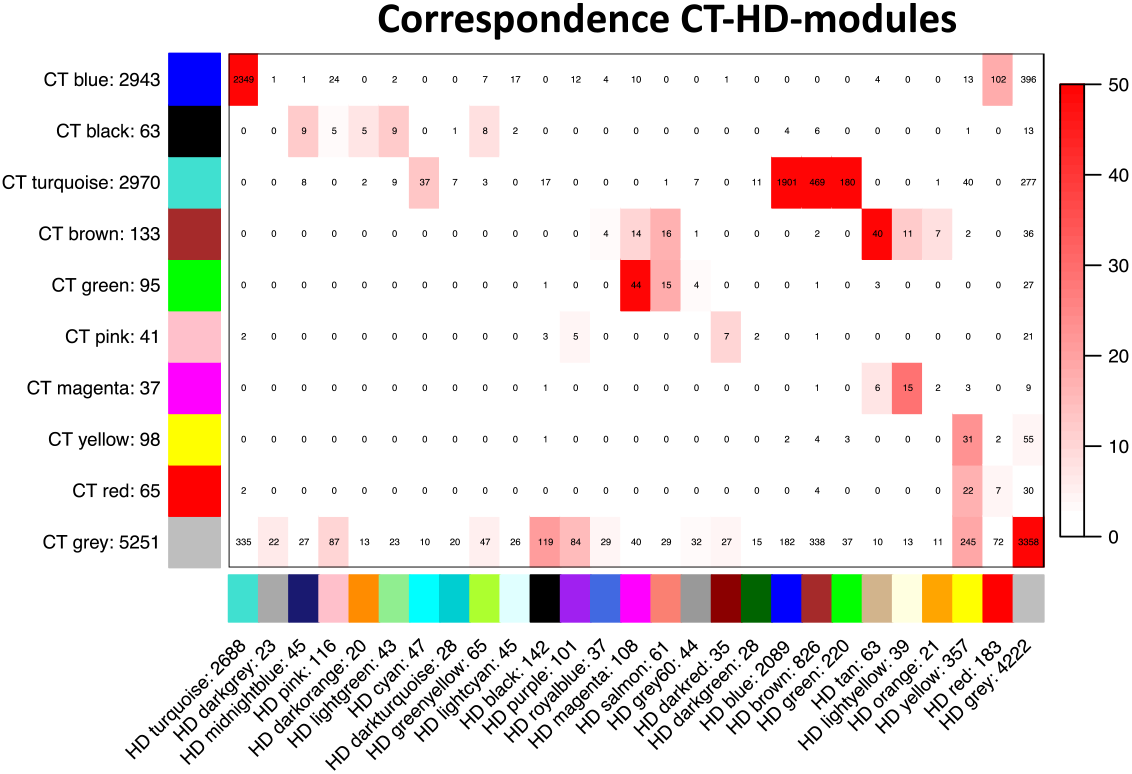
Module overlap between gene co-expression networks of CT lines and HD lines at generation 1. The number of genes in each module and their overlap is shown. The intensity of red indicates significance of overlap tested with Fisher’s exact test (-log10(p-value)).

**Supplementary figure 4:**
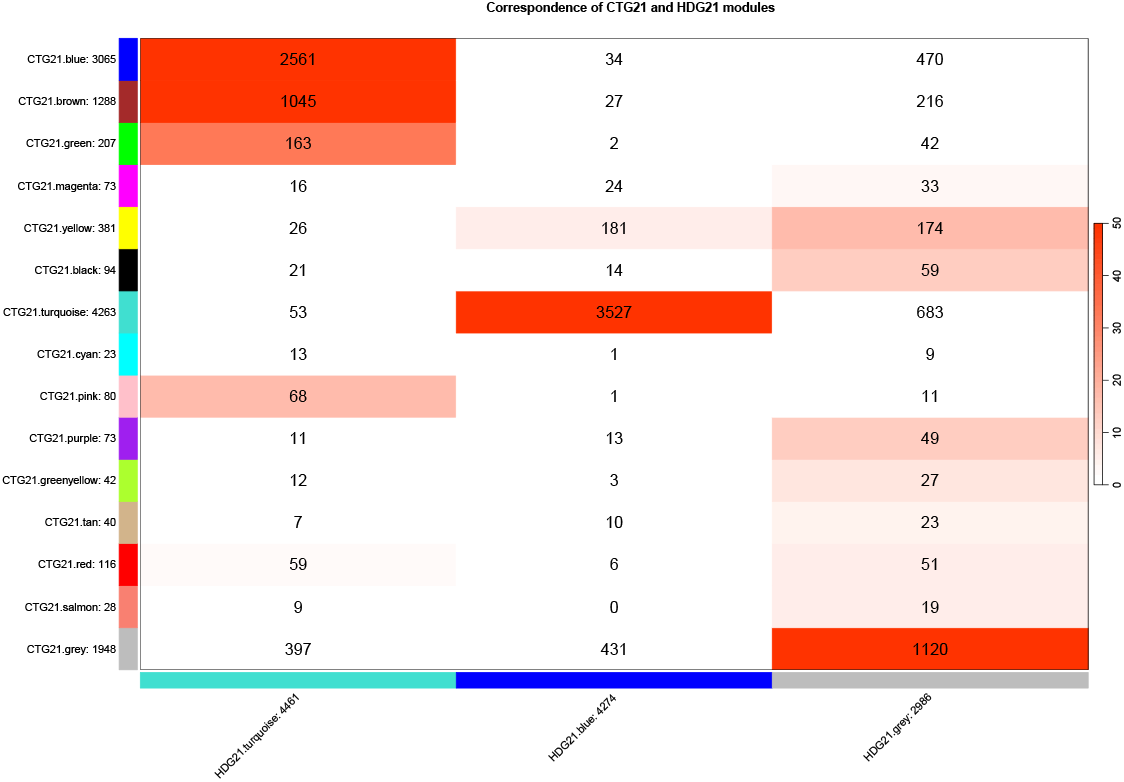
Module overlap between gene co-expression networks of CT lines and HD lines at generation 22. The number of genes in each module and their overlap is shown. The intensity of red indicates significance of overlap tested with Fisher’s exact test (-log10(p-value)).

**Supplementary figure 5:**
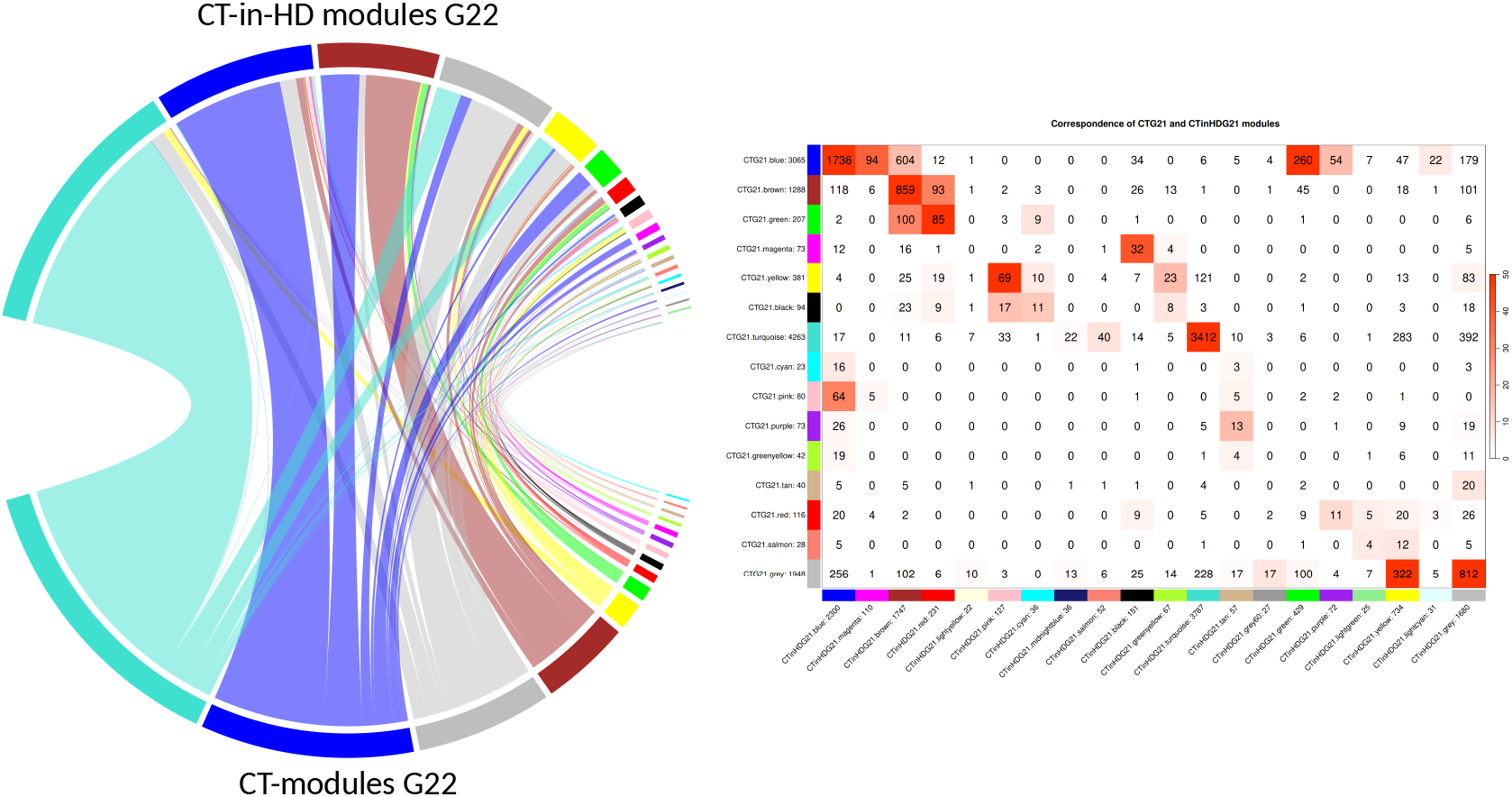
Module overlap between gene co-expression networks in CT lines and CT lines who were transplanted in HD at generation 22. The number of genes in each module and their overlap is shown in the table on the right. The intensity of red indicates significance of overlap tested with Fisher’s exact test (-log10(p-value)).

**Supplementary figure 6:**
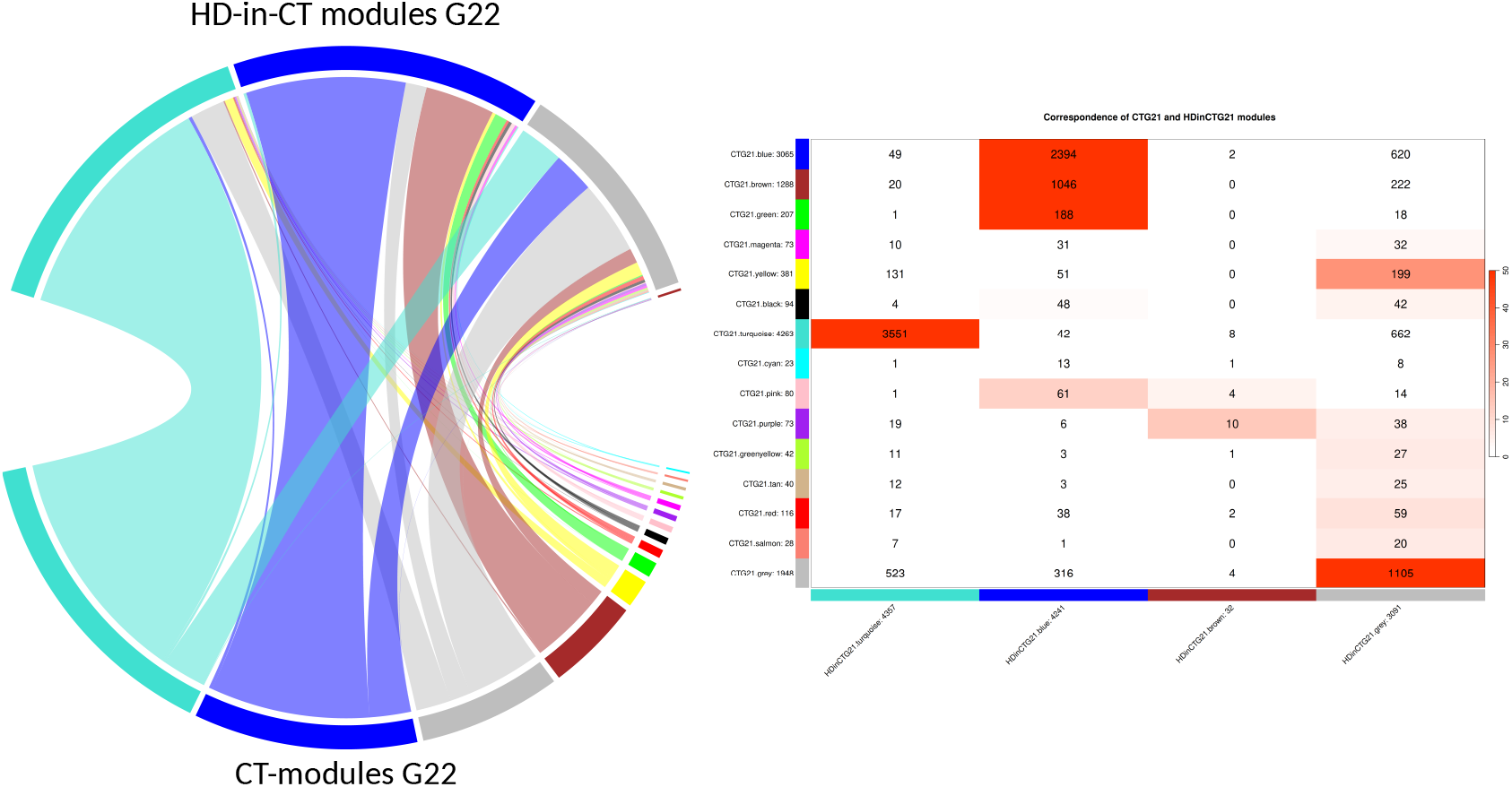
Module overlap between gene co-expression networks of CT lines and HD lines who were transplanted in CT at generation 22. The number of genes in each module and their overlap is shown in the table on the right. The intensity of red indicates significance of overlap tested with Fisher’s exact test (-log10(p-value)).

**Supplementary figure 7:**
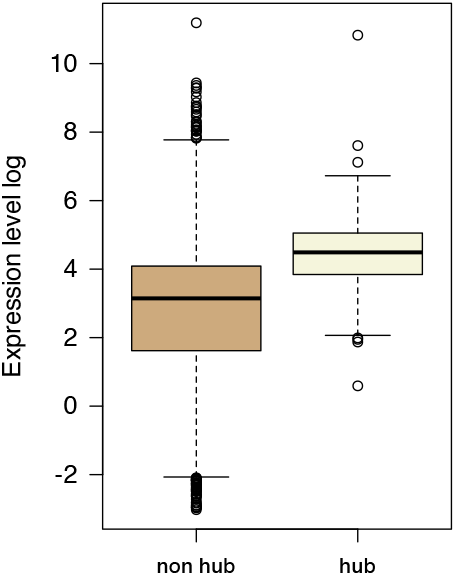
Expression levels of non-hub and hub genes (defined as the top ten genes of each model showing the highest intramodular connectivity). Expression levels were corrected for batch effects (sequencing runs) and mother identity to account for potential maternal and common environmental effects.

**Supplementary figure 8:**
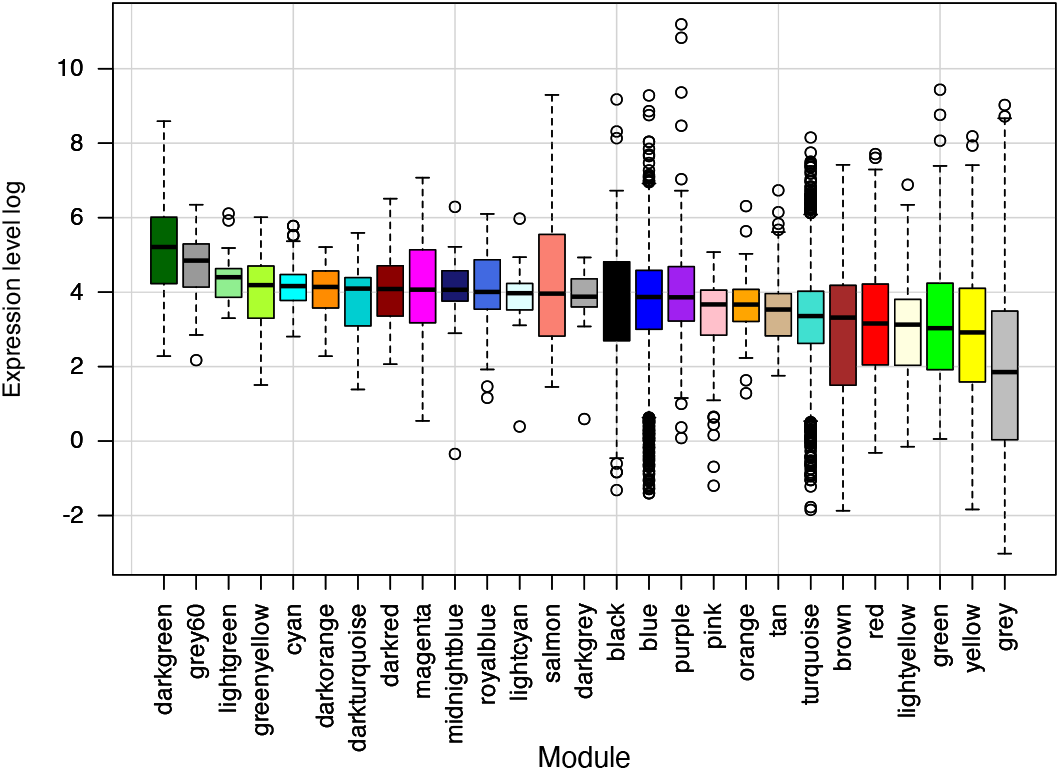
Expression levels of genes assigned to different modules in hot-dry (G1). The grey module contains unassigned genes. Expression levels were corrected for sequencing runs and mother identity.

**Supplementary figure 9:**
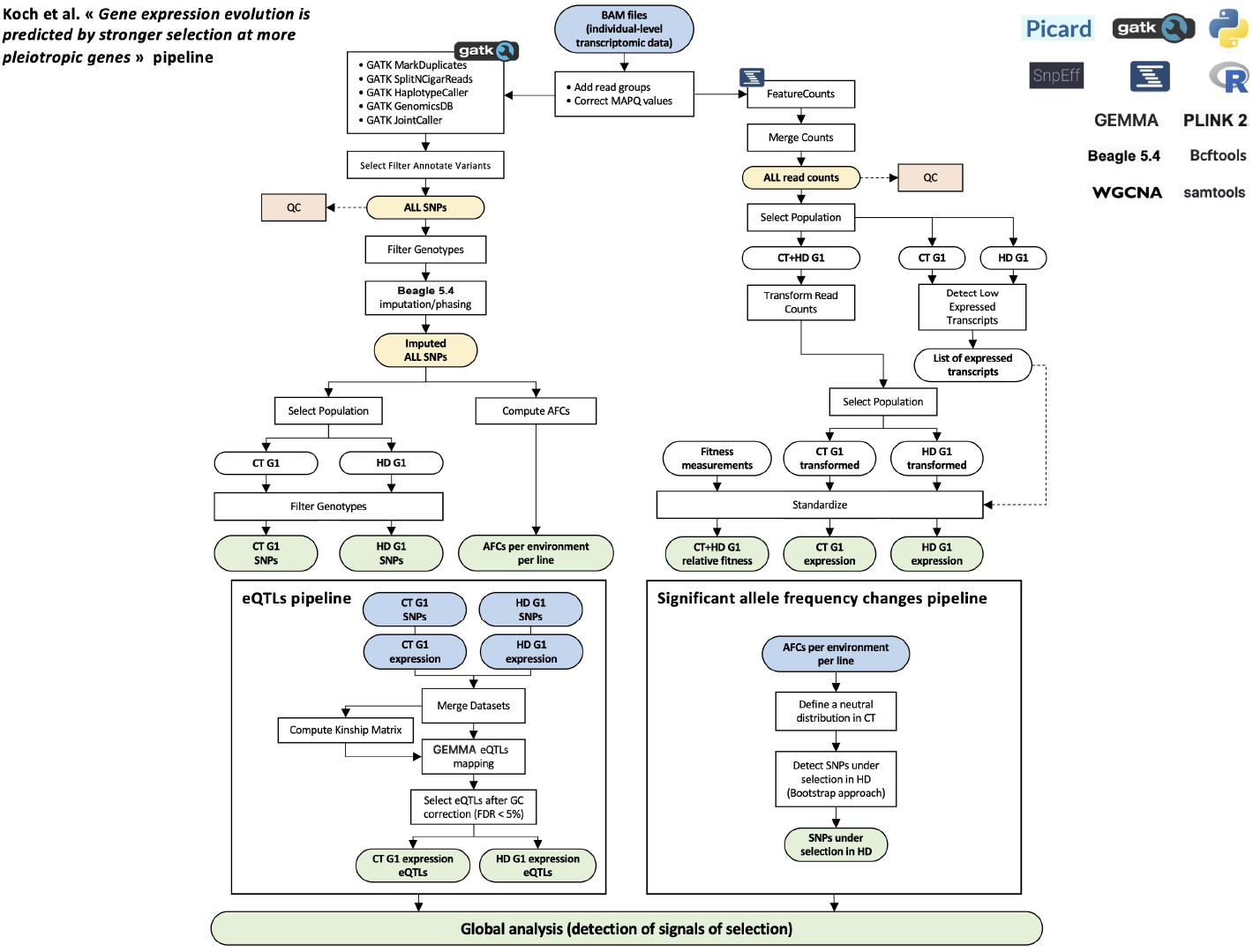
Overview of the bioinformatics pipelines. The bioinformatics pipelines used to recover single nucleotide polymorphism, eQTLs and allele frequency changes (AFCs) are summarized in this diagram.

**Supplementary figure 10:**
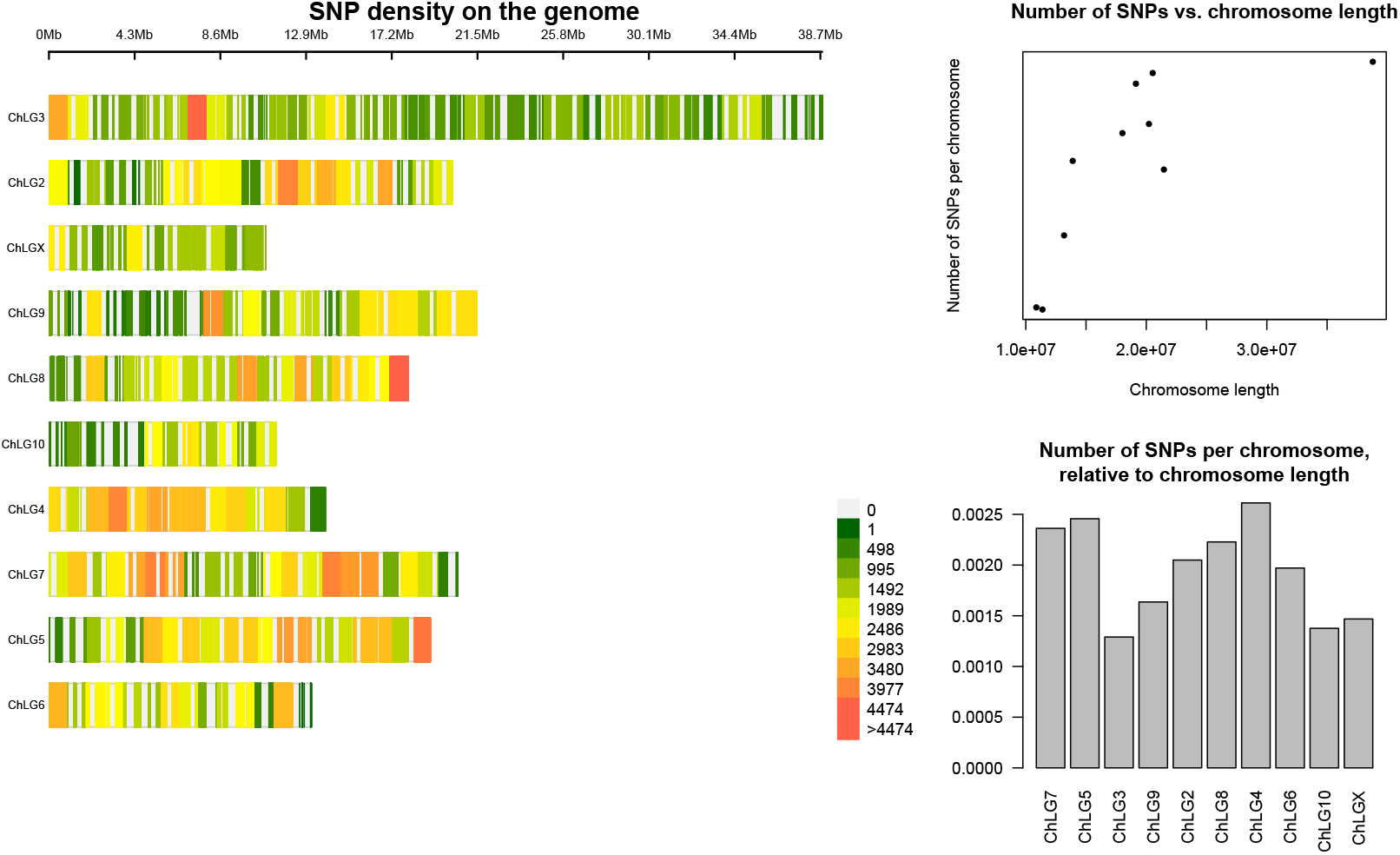
Repartition of SNPs on the genome. **Left panel:** SNP density along the genome. **Top-right panel:** Number of SNPs per chromosome vs. chromosome length for the 10 chromosomes of *Tribolium castaneum*. **Bottom-right panel**: Number of SNPs per chromosome relative to chromosome length.

**Supplementary figure 11:**
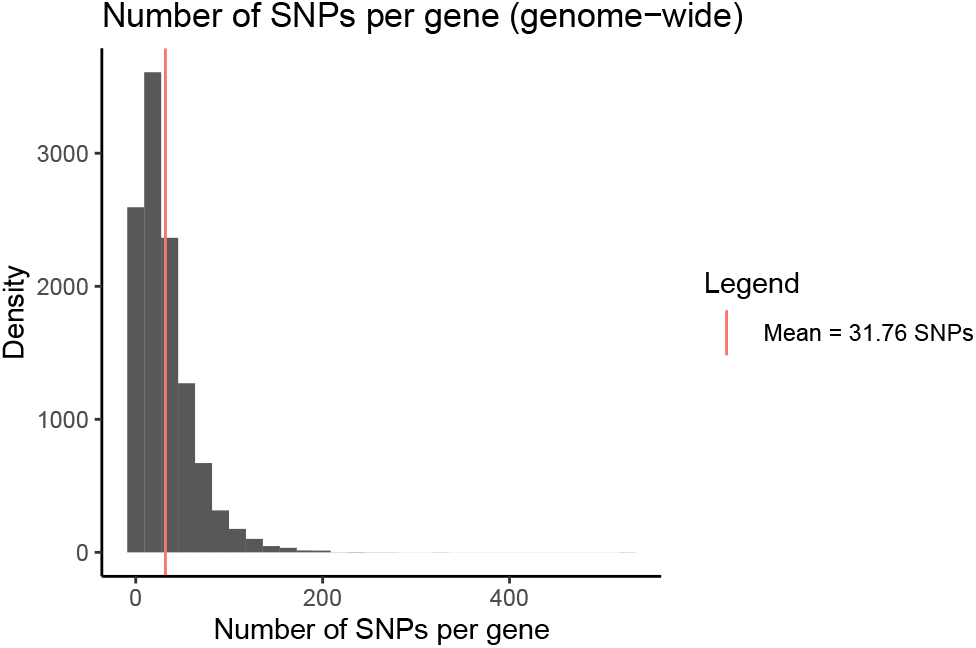
Number of SNPs per gene in the genome. Histogram of the number of SNPs per gene on the entire genome. The mean number of SNPs per gene is indicated by a red vertical line (31.76 SNPs).

**Supplementary figure 12:**
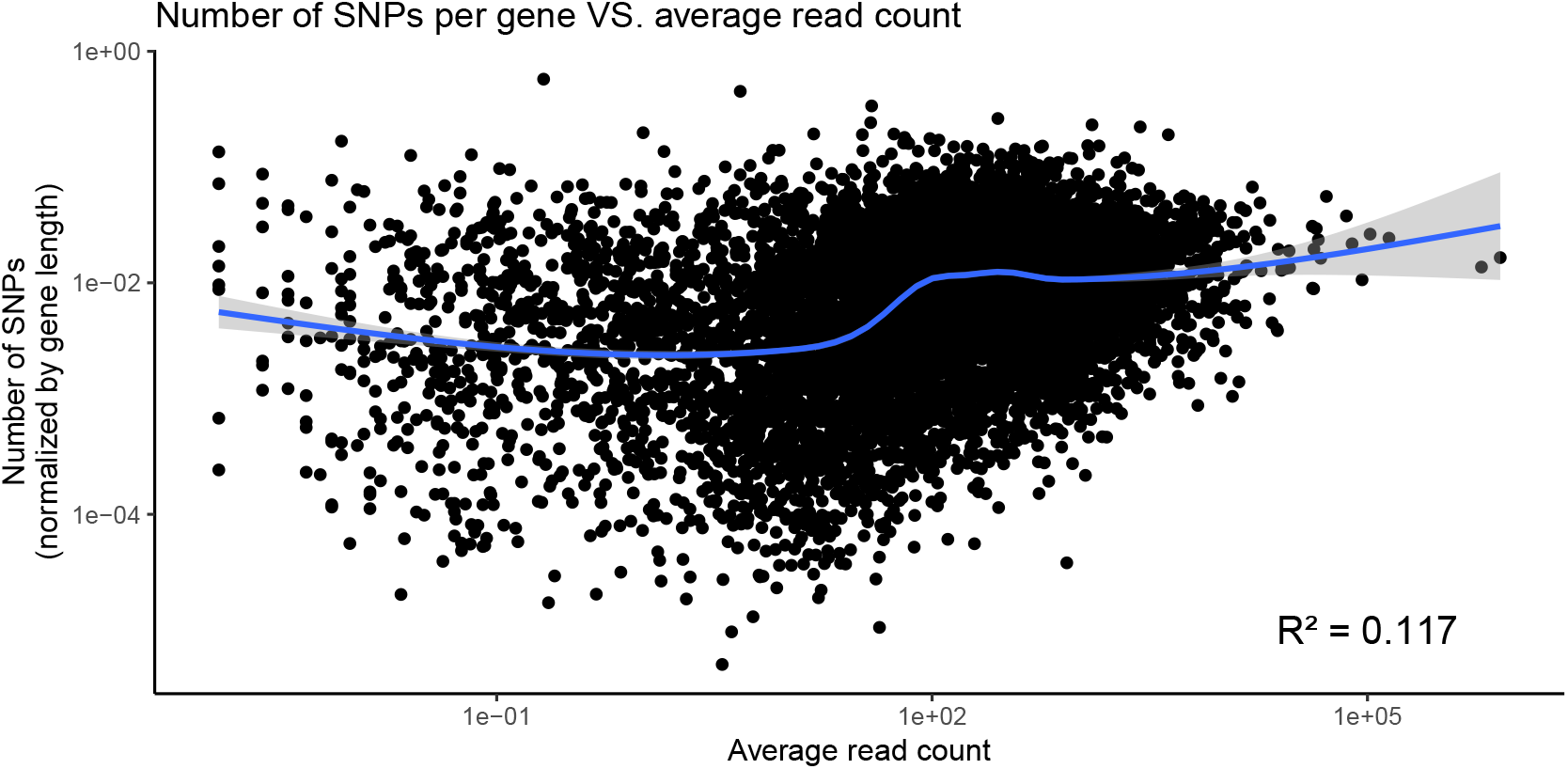
Number of detected SNPs per gene versus their average read count. The number of detected SNPs per gene (normalized per gene length) is compared to the average read count per gene, across all sequenced individuals. The read count data is used before transformation and normalization to allow for the comparison of expression levels. The blue solid line and grey shaded area show a smoothed curve and its confidence interval. Linear regression *R*^2^ = 0.117.

**Supplementary figure 13:**
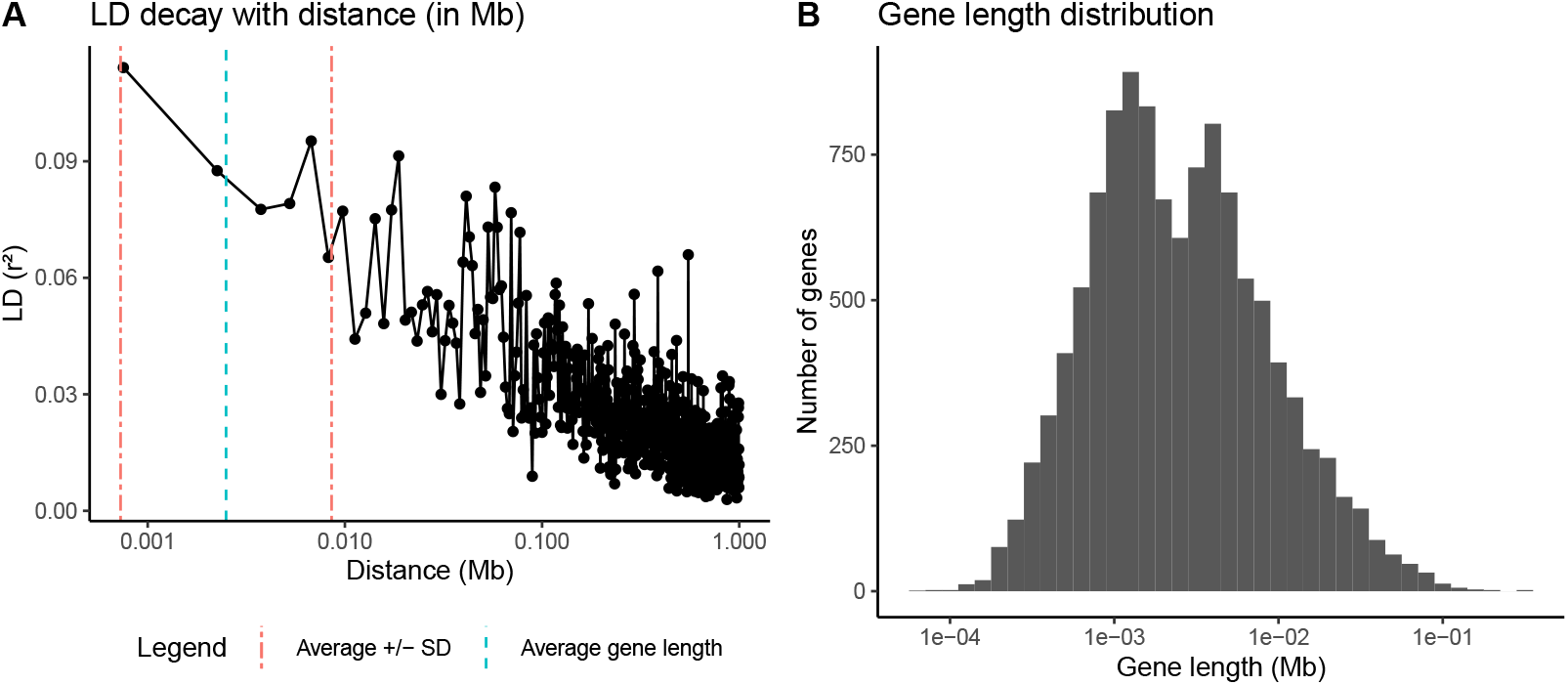
Linkage disequilibrium decay with distance. **(A)** LD decay (*r*^2^) with distance (in Mb). Windows of 1.5Kb were used to average the LD. The blue dashed line indicates the average length of a gene (∼2.5Kb). The red dashed lines indicates the gene length average +/-the standard deviation. **(B)** Distribution of gene length in log-scale. Linkage disequilibrium (LD) decay was calculated on each of the 10 chromosome applying PLINK1.9 software (Chang et al., 2015) on the final VCF file (358,142 SNPs). Variants with a minor allele frequency < 0.05 were removed (–maf 0.05), individuals with more than 50% of missing genotypes were also filtered out (–mind 0.5). For LD calculation, the minimum distance between two SNPs was 100bp (–ld-window 100) and the maximum window for calculation was 1Mb (-ld-window-kb 1000). The VCF file was also thinned by only keeping 1% of SNPs (–thin 0.01), in order to facilitate further calculations.

**Supplementary figure 14:**
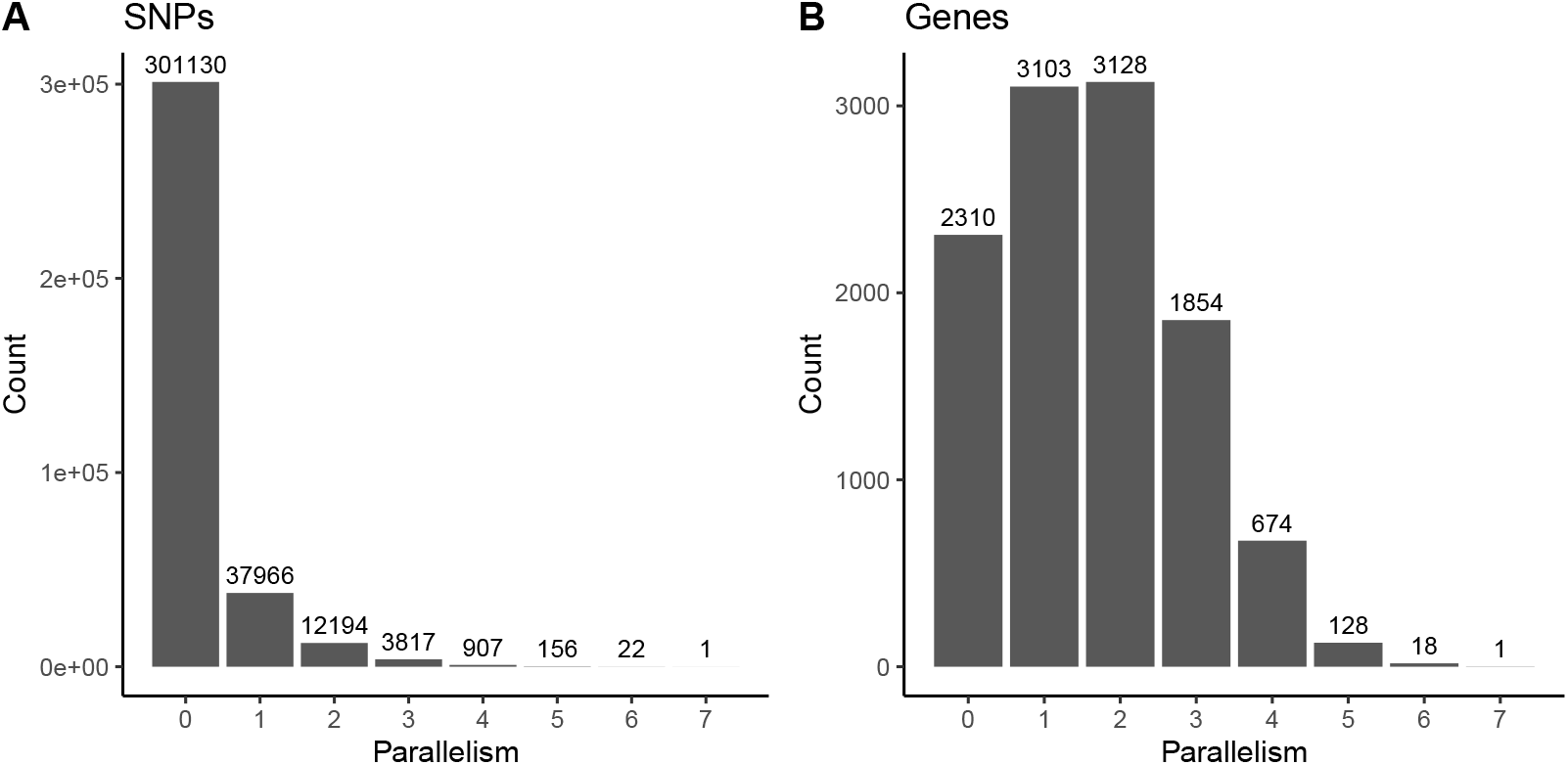
Parallelism of SNPs and genes frequency shifts. Distribution of the parallelism of frequency shifts, from no shift (0) to fully parallel shifts in the seven lines for (**A**) SNPs, (**B**) genes. The count is shown in each category.

**Supplementary figure 15:**
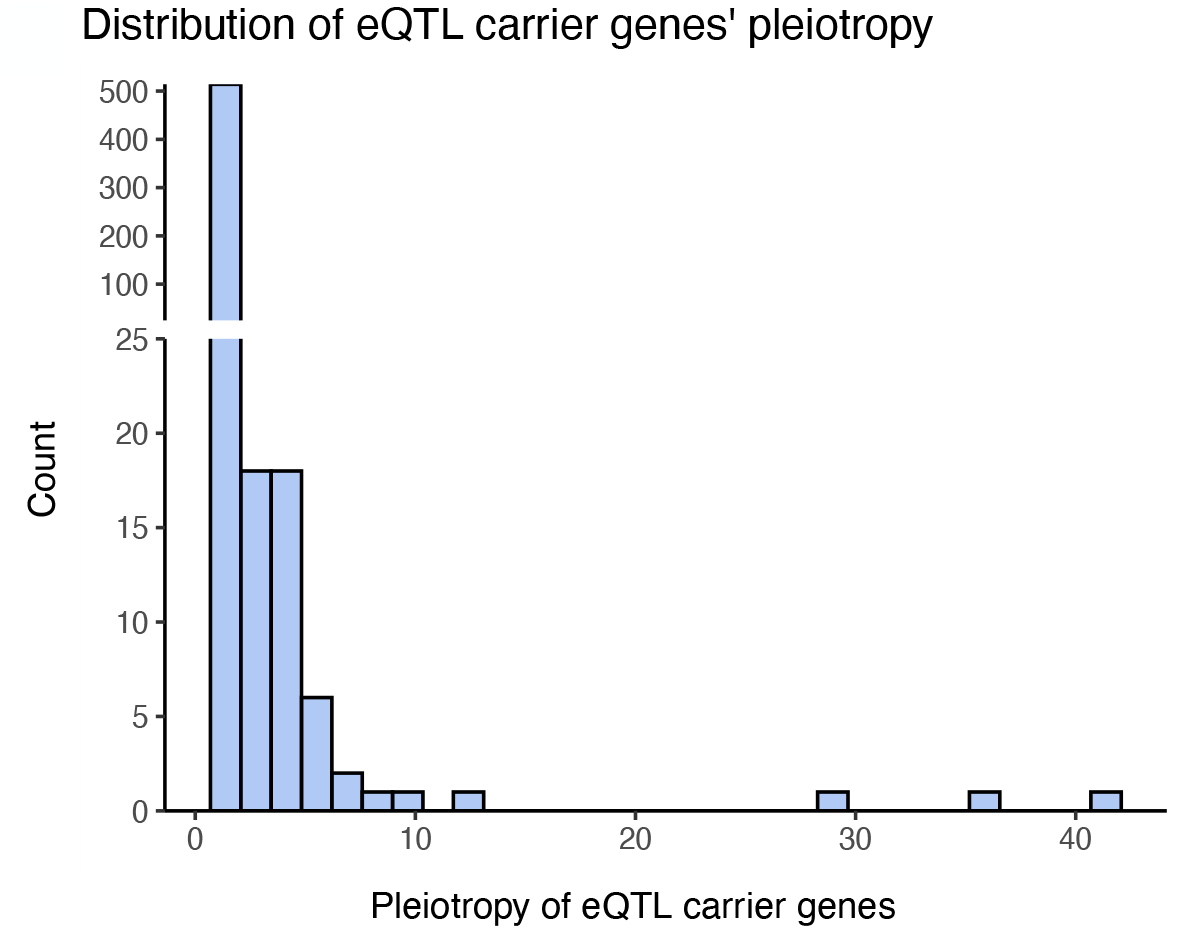
Distributions of the pleiotropy of eQTL carrier genes. Distribution of the pleiotropy (*i*.*e*., the number of significantly associated phenotypes) for all 564 eQTL carrier genes. Pleiotropy values range from 1 to 41. Vertical axis was re-scaled above *y* = 25 for readability.

## Supporting data online

The following scripts and data sets are accessible online at https://doi.org/10.5281/zenodo.15298510.

**Script S1**. BSFG estimates (50.9 MB).

**Script S2**. WGCNA analysis and results G1 (99.6 MB).

**Data S1**. Datasets resulting from the global genomics analysis of the output of the transcriptomics pipeline (5.9 MB).

**Data S2**. VCF file containing the 566,296 quality-checked SNPs (1.1 GB).

**Data S3**. VCF file containing the 358,142 SNPs with a call rate higher or equal to 50% (964.3 MB).

**Data S4**. VCF file containing the imputed genotypes (964.3 MB).

**Data S5**. Results of the imputation tests (479.5 kB).

**Data S6**. VCF file containing imputed genotypes where SNPs with a minor allele frequency lower than 0.05 have been filtered out (MAF ≤ 0.05) (176.0 MB).

**Data S7**. List of the 1,273 significant eQTL associations (for gene expression levels or relative fitness as phenotypes) (61.2 kB).

**Data S8**. List of allele frequency changes (AFCs) for every markers in HD environment, for lines L1, L2, L3, L5, L6, Mx1 and Mx2 (32.4 MB).

**Data S9**. List of all SNPs indicating if their AFC is significantly higher in each line and their degree of parallelism (2.4 MB).

**Data S10**. Excel file containing the results of the gene functional enrichment analysis of the hub and eQTL carrier genes (33.7 KB).

